# Large differences in photorespiration and its temperature response among temperate trees

**DOI:** 10.1101/2025.11.22.689893

**Authors:** Rakesh Tiwari, Philip David, Robert Muscarella

**Author notes:** The full-length article is available in Kannada and Sanskrit languages in the Supporting section S1 and S2.

## Abstract

Photorespiration significantly influences terrestrial carbon fluxes, yet empirical measurements of its variability across tree species and temperature conditions remain limited, constraining predictions of vegetation and climate models. We quantified apparent photorespiratory CO_2_ loss (*L*_app_) and its temperature response for seven temperate broadleaf tree species in northern Europe, using *in situ* O_2_-shift measurements in Uppsala, Sweden during peak summer. Apparent loss was derived as the difference between net CO_2_ assimilation under ambient (*A*_net_) and O_2_-free conditions at three leaf temperatures (25, 30, and 35 °C), spanning typical and heat-wave scenarios. Apparent photorespiratory CO_2_ loss showed pronounced interspecific variation and increased with temperature, while net photosynthesis remained relatively stable. The ratio of apparent loss to net photosynthesis (ϕ = *L*_app_/*A*_net_) rose sharply with temperature, reaching species-mean values up to 0.94 at 35 °C, indicating that photorespiration can represent nearly the entirety of net carbon gain under heat stress even when leaves remain net CO_2_ sinks. Suppression of photorespiration under N_2_ and associated changes in leaf temperature systematically reallocated photosynthetic electron transport: the fraction of ambient electron transport rate (ETR) allocated to net CO_2_ assimilation declined with temperature, whereas the complementary fraction allocated to apparent photorespiratory loss and other O_2_-dependent sinks increased, with ETR-based apparent loss and its proportional expression rising steeply across the 25–35 °C range. Together, these *in situ* flux and partitioning measurements reveal high variability and strong temperature sensitivity in apparent photorespiration among temperate trees. Compared to crop-based parameterisations, the ϕ values we report for temperate trees are substantially higher and more temperature-dependent, providing species-specific constraints that can improve Farquhar–von Caemmerer–Berry–type vegetation model representations of photorespiration in forest ecosystems.

## Introduction

Photorespiration, the second largest biological flux on Earth, has major impacts on terrestrial carbon fluxes but we have limited understanding of how this process varies across environmental conditions and among species (Bauwe *et al*., 2010; Busch & Sage, 2017). The photorespiration process occurs when Rubisco, the key enzyme responsible for catalysing photosynthesis by binding CO_2_ to its substrates, catalyses the binding of O_2_ (Lorimer & Andrews, 1973). This oxygenation reaction produces glycolate (2-phosphoglycolate), a toxic intermediate that is subsequently recycled through the photorespiratory pathway (Ogren & Bowes, 1971). During this process, a portion of the previously assimilated carbon is released as CO_2_, generating substantial photorespiratory CO_2_ loss at the leaf level (Wingler *et al*., 2000). Despite the large impact of photorespiration on overall photosynthetic yield and the global carbon cycle, empirical studies quantifying variation in photorespiration CO_2_ loss at the leaf level remain scarce.

Most dynamic global vegetation models (DGVMs) integrate photorespiration theoretically through the Farquhar–von Caemmerer–Berry (FvCB) model for C3 plants (Farquhar *et al*., 1980). In part due to a lack of empirical data, this approach does not explicitly incorporate variation of photorespiration rates across species or climate conditions (Beauclaire et al., 2024; Oberpriller et al., 2022). In practice, DGVMs that use the FvCB model typically adopt a fixed value for the ratio of photorespiration (*R*_p_) to photosynthesis (*A*_net_) – expressed as ϕ = *R*_p_/*A*_net_ (Box 1). Notably, the empirical basis for the ϕ parameter relies primarily on crop plant data - namely rice, wheat, and soybean - with only a limited number of studies on trees (Epron et al., 1995; Hochberg et al., 2013; Huang et al., 2015; Keys et al., 1977; Tomeo & Rosenthal, 2018; Valentini et al., 1995; Z.-P. Ye et al., 2019; Yeo et al., 1994; C. Zhang et al., 2016). Consequently, the FvCB model may inadequately represent photorespiration in forest ecosystems and lead to inaccurate estimations of other critical parameters (e.g., maximum rubisco carboxylation rate and electron transport rates – V_cmax_ and *J*_max_), thereby reducing accuracy of vegetation–climate feedback projections (Lochocki & McGrath, 2025; Z. Ye et al., 2025; Z.-P. Ye et al., 2025). Empirical constraints, particularly for long-lived plants, could help improve model fidelity and provide key insights to the role of photorespiration in the global carbon cycle (van Bodegom *et al*., 2014).

Photorespiration consumes a substantial fraction of the chemical energy generated during the light reactions of photosynthesis, thereby reducing photosynthetic light-use efficiency and has thus been considered a major inefficiency in plant productivity (Zhu *et al*., 2010). However, under stress conditions when photosynthesis is suppressed (e.g., high temperature), photorespiration serves as an essential alternative sink for the light reactions products (Huang et al., 2019; Osei-Bonsu et al., 2021; Streb et al., 2005), alleviates photoinhibition of photosystem I (Shi et al., 2022; Wada et al., 2020), and prevents the accumulation of reactive oxygen species (Foyer *et al*., 1994, 2009; Jiang *et al*., 2023), thereby providing thermal protection (A. P. Cavanagh et al., 2022; Chuang & Ling, 2005; Hu et al., 2020; Kozaki & Takeba, 1996; Wingler et al., 2000). The protective role of photorespiration is supported by studies on *Arabidopsis*, tobacco, rice, and maize mutants deficient in photorespiratory enzymes (Timm et al., 2012; Zelitch et al., 2009). The loss of photorespiratory function in these mutants led to increased heat sensitivity, impaired growth, leaf necrosis, and reduced survival. Thus, photorespiration is recognized as a key component of plant thermal tolerance, maintaining redox and energy balance and supporting cellular homeostasis during thermal stress (Bauwe et al., 2010; Noctor et al., 2002; Roze et al., 2025; Voss et al., 2013). However, our knowledge of the temperature sensitivity of photorespiration remains limited, particularly for perennial plants.

Variation among tree species in terms of photorespiration rates may be related to other aspects of life-history. For example, photosynthetic capacity has been shown to vary systematically across successional stages: early-successional species tend to exhibit higher net photosynthesis and Rubisco carboxylation rates, prioritizing rapid carbon acquisition, whereas late-successional species adopt more conservative metabolic strategies (G. Zhang et al., 2018; Ziegler et al., 2020). Thermal responses also differ markedly, with early-successional trees maintaining stable performance under warming while late-successional species tend to show greater heat vulnerability (Mujawamariya *et al*., 2023). Given the critical role of photorespiration in thermal protection, it remains unclear whether species with higher photosynthetic rates also exhibit proportionally higher photorespiration or if these processes are decoupled, particularly under thermal stress.

### Box 1

Values for the ratio of photorespiration (*R*_p_) and photosynthesis (*A*_net_) rates. Letter annotations indicate significant differences between means of the groups.

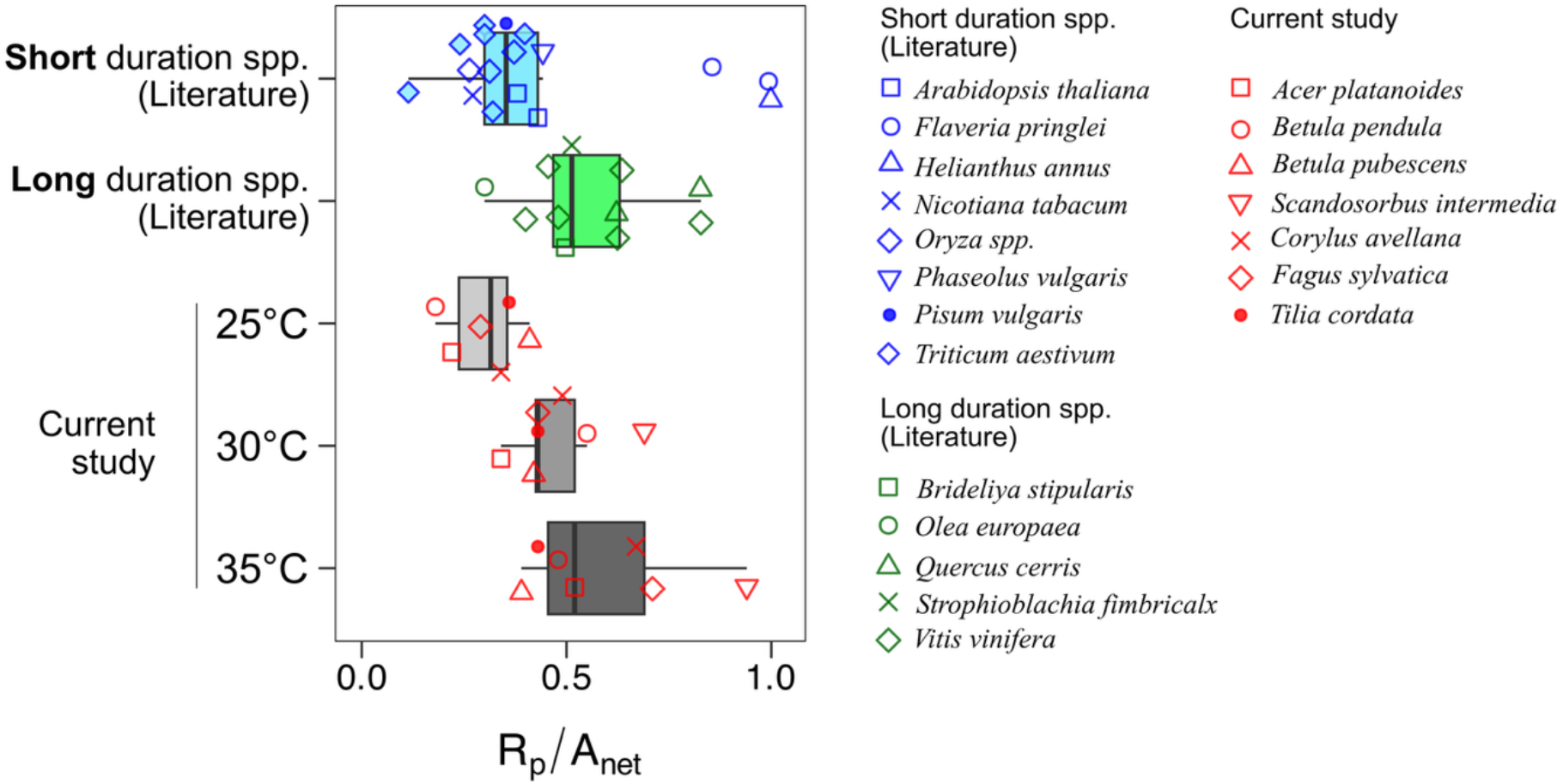

Compared with photosynthesis (*A*_net_), *in situ*, leaf-level measurements of photorespiratory CO_2_ release (*R*_p_) rates are relatively rare and have been reported for only a limited number of species. Two main approaches are commonly used to estimate *R*_p_: (a) comparing photosynthetic CO_2_ assimilation rates under ambient and low-oxygen conditions to suppress photorespiration (Sharkey, 1985, 1988; Sharkey *et al*., 1988), and (b) using simultaneous measurements of photosynthetic electron transport rates with a fixed partitioning ratio between photorespiratory and non-photorespiratory pathways (Peterson, 1989; Valentini *et al*., 1995; Yoshimura *et al*., 2001). Both methods have notable limitations and require corrections for alternative CO_2_ release pathways (Busch, 2013; Busch et al., 2017; Sharkey, 1988; Xu et al., 2021). The figure above compiles data from both approaches, including only studies that reported *R*_p_ and *A*_net_ with sufficient information on corrections for alternative pathway CO_2_ release (Supplementary Table S1); these compiled literature values are therefore corrected estimates of biochemical *R*_p_. From these data, we calculated the ratio of *R*_p_ to *A*_net_ (ϕ) at a leaf temperature of 25°C, highlighting the variation in ϕ values across species. The data is separated based on short duration species and long duration species that include trees and woody perennials. Species average values from the current study are also presented at each of three temperature points; because our estimates are derived from an uncorrected O_2_-shift approach, they represent apparent photorespiratory CO_2_ loss rather than a fully partitioned biochemical *R*_p_ and likely reflect an upper-bound approximation given residual mitochondrial respiration and internally generated O_2_ under N_2_ (see Methods section). This distinction between corrected literature *R*_p_ values and our uncorrected apparent estimates should be considered when interpreting absolute magnitudes across the two datasets, though relative patterns across species and temperature remain informative.

Beyond Rubisco kinetics, heat stress introduces several factors that may disrupt the coupling between photosynthesis and photorespiration. At temperatures above ∼42 °C, PSII reaction centre proteins rapidly dephosphorylate and destabilize, reducing electron transport and increasing photoinhibition (Rokka *et al*., 2000; Salvucci & Crafts-Brandner, 2004). Stomatal conductance may remain high or even rise under heat to enhance transpiration cooling, yet net photosynthesis often declines due to biochemical limitations, resulting in a decoupling between stomatal behaviour and carbon assimilation (Diao et al., 2024; Marchin et al., 2023). Additionally, heat-driven reactive oxygen species production activates antioxidant defences that preferentially support photorespiratory pathways (Kuppusamy *et al*., 2023). Consequently, under higher temperatures, the shifting metabolic and energetic costs of photosynthesis and photorespiration is likely affecting their relative rates. In many cases, heat stress alters stomatal behaviour and internal CO_2_ supply only modestly, while biochemical limitations and photorespiratory flux dominate changes in net assimilation efficiency (Diao et al., 2024; Marchin et al., 2023).

Recent syntheses have highlighted that photorespiration may consume 25%–50% of gross CO_2_ assimilation in C3 leaves under thermal stress and that tree-specific temperature responses and electron transport partitioning remain critically undermeasured (Tiwari *et al*., 2026). Here, we explicitly quantify apparent photorespiratory CO_2_ loss and the partitioning of photosynthetic electron transport between net CO_2_ assimilation and apparent photorespiration across seven temperate broadleaf tree species and three leaf temperatures, providing *in situ* constraints on how warming redistributes electron use in forest trees.

Electron transport rate (ETR) partitioning provides an independent, complementary approach to O_2_-shift methods for probing photorespiratory flux: because photorespiration consumes electrons through downstream glycolate metabolism, the allocation of ambient ETR between net CO_2_ assimilation and photorespiratory pathways can be estimated by comparing ETR under ambient and low-oxygen conditions (Peterson, 1989; Valentini et al., 1995; Yoshimura et al., 2001). However, this approach carries its own uncertainties, as changes in ETR may partly reflect alternative electron sinks such as cyclic electron flow or the Mehler reaction rather than photorespiration alone (Takagi et al., 2016), and it has rarely been applied or validated in woody species (Tiwari et al., 2026). Combining this electron-transport-based partitioning with direct O_2_-shift measurements of *A*_net_ therefore allows two independently derived, imperfect estimates of photorespiratory response to be cross-checked against one another, strengthening inference beyond either approach alone.

We selected a set of seven widely distributed broadleaf tree species from the northern temperate region and conducted *in situ* measurements of *R*_p_ and *A*_net_ at three leaf temperatures: 25°C (typical summer conditions in the study area), 30°C (occasional summer highs), and 35°C (heat wave conditions, which are increasingly common in the study area). From these measurements, we derived apparent photorespiratory CO_2_ loss as the difference between net assimilation under O_2_-free and ambient conditions. We addressed the following questions: (1) How does the magnitude of apparent photorespiratory CO_2_ loss (*L*_app_) vary among species? (2) How does apparent photorespiratory CO_2_ loss (ϕ = *L*_app_/*A*_net_) respond to temperature? (3) How is photosynthetic electron transport (ETR) partitioned between net CO_2_ assimilation and apparent photorespiratory CO_2_ loss, and how does this partitioning vary among species and with temperature?

Based on existing studies, we hypothesized that net photosynthesis would remain relatively stable or decline modestly across our temperature range, while apparent photorespiratory CO_2_ loss would increase with temperature due to the temperature sensitivity of Rubisco oxygenation and associated CO_2_ release. We further hypothesized that the proportional carbon cost of photorespiration, expressed as ϕ = *L*_app_/*A*_net_, would increase with warming and reach high values under heat-wave conditions (35°C). Finally, we expected electron transport partitioning to shift progressively away from net CO_2_ assimilation and towards apparent photorespiratory loss at elevated leaf temperatures (Tomeo & Rosenthal, 2018; B. J. Walker et al., 2017).

## Materials and Methods

### Study site

We conducted this study during the summer of 2024 (June–August) in Stadsskogen nature reserve, Uppsala, Sweden (59.842563N, 17.621133E; 40 m a.s.l.). The site experiences a temperate continental climate, with an average annual temperature of approximately 7 °C and average annual precipitation of 576 mm. During the study period, daytime temperatures (8-17h) ranged from 4.2 °C to 33.3 °C, with a mean of 20.1 °C. Notably, the area holds the Swedish record for the highest observed summer temperature, reaching 38 °C in nearby Ultuna.

### Study species

Measurements were conducted on seven common broadleaf tree species: silver birch (*Betula pendula* Roth.), downy birch (*Betula pubescens* Ehrh.) European beech (*Fagus sylvatica* L.), Swedish whitebeam (*Scandosorbus intermedia* (Ehrh.) Sennikov), Norway maple (*Acer platanoides* L.), small-leaved linden (*Tilia cordata* Mill.) and common hazel (*Corylus avellana* L.). The study species encompass a large portion of the common non-conifer trees found in the deciduous forests of Sweden and most are common across Europe (Rydin *et al*., 1999). The study area represents the northern range edge of *F. sylvatica, S. intermedia*, *A. platanoides*, *T. cordata*, and *C. avellana*, whereas it represents a more central part of the geographic distribution for the *B. pendula* and *B. pubescens* (de Rigo *et al*., 2016).

The seven study species represent a range of life-histories, with distinct resource acquisition strategies and successional associations. *B. pendula* is an early successional species with an acquisitive resource strategy characterized by high photosynthetic rates under high light conditions and rapid biomass growth rates (Õunapuu-Pikas *et al*., 2025). Five species, *B. pubescens*, *A. platanoides*, *T. cordata*, *C. avellana* and *S. intermedia* are more typically associated with mid-successional stages, exhibiting intermediate shade tolerance and demonstrating flexible resource strategies that allow rapid response to canopy gaps while maintaining modest performance under moderate shade conditions (Hagemeier & Leuschner, 2019a). *F. sylvatica* represents a typical late-successional species with conservative resource strategies, featuring shade-tolerance mechanisms including high specific leaf area plasticity, efficient light capture at photon flux densities as low as 1-3% of full sunlight, and high competitive ability through efficient shade production that reduces understory light availability (Desotgiu *et al*., 2012; Hagemeier & Leuschner, 2019b; Leuschner & Hagemeier, 2020).

### Gas exchange measurements

We measured light-saturated net CO_2_ assimilation rate (*A*_net_) using a Li-6800 infrared gas analyser (IRGA) with a Li-6800-01A leaf chamber (Li-Cor, Lincoln, Nebraska). Leaf environmental conditions included a CO_2_ concentration of 420 μ mol mol⁻¹, 50% relative humidity, and an irradiance of 1500 μ mol m⁻² s⁻¹. Steady-state measurements were conducted on three fully mature, healthy leaves in the upper canopy strata for at least five individual trees per species. We made measurements at three leaf temperatures: 25 °C (typical summer conditions in Uppsala), 30 °C (occasional summer highs), and 35 °C (heat-wave conditions, now increasingly common in the region) (SMHI, 2024) (see Supplementary Figure S1). Measurements were taken between 10:00 and 16:00. For each leaf, net CO_2_ assimilation (*A*_net_, referred to as *A*) was first measured under ambient O_2_ (21% O_2_) to record photosynthesis including photorespiration (*A*_net,_ ambient), and then under an O_2_-free nitrogen atmosphere to record *A*_net_ with photorespiration strongly suppressed (***A***_***N***2_). To achieve an O_2_-free environment, we used nitrogen gas (99.8% purity; O_2_ < 0.01%, Air Liquide) as IRGA input at a flow rate of ∼1.6 L min⁻¹. Leaves were exposed to the N_2_ atmosphere for 1–5 min prior to measurement to allow chamber and leaf gas composition to stabilise and O_2_ to be flushed from the system. The transition duration varied with the rate of chamber flushing and leaf gas exchange, and in all cases the criterion for proceeding was the attainment of a new CO2 flux steady state, confirmed from the LI-6800 real-time trace before logging. Upon reaching steady state, we recorded *A*_net_ under these nominally non-photorespiratory conditions as ***A***_***N***2_. Before subsequent measurements, we flushed the chamber with ambient air and allowed CO_2_ flux from the leaf to re-stabilise under the desired conditions.

In addition to *A*_net_, we recorded stomatal conductance to water vapor (*g*_s_) under both ambient O_2_ and N_2_ conditions for each measurement pair, as an output of the LI-6800 gas exchange system, to assess whether the O_2_ shift affected stomatal response in addition to biochemical assimilation. Similarly, we recorded chlorophyll fluorescence and the instrument-derived estimate of linear electron transport rate (ETR) for each measurement under both ambient O_2_ and N_2_, using the default LI-6800 fluorescence protocol. These ETR measurements were used to quantify how suppression of photorespiration and changes in leaf temperature affected photosynthetic electron transport, and to partition ambient ETR between net CO_2_ assimilation and apparent photorespiratory CO_2_ loss using paired ambient–N_2_ measurements (see “ETR partitioning calculations” below).

Explicitly quantifying stomatal conductance and the ratio of CO_2_ assimilation to intercellular CO_2_ concentration (*A*_net_/*C*_i_) alongside apparent photorespiratory CO_2_ loss is important to assess whether warming-induced changes in net photosynthesis primarily reflect stomatal or biochemical controls.

### Apparent photorespiratory CO2 loss calculations

We quantified apparent photorespiratory CO_2_ loss (hereafter “apparent loss”) by comparing net CO_2_ assimilation under ambient and O_2_-free conditions for each leaf. For a given measurement pair, we defined apparent loss as

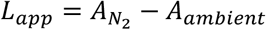

where *A*_ambient_ is net CO_2_ assimilation measured at 21% O_2_ and ***A***_***N***2_ is net CO_2_ assimilation measured under the N_2_ atmosphere at otherwise identical conditions. Positive values of ***L***_***app***_ thus indicate that suppression of photorespiration increases net CO_2_ assimilation, consistent with photorespiration being a net carbon cost under our conditions. To compare across species and temperatures, we also calculated the ratio of apparent loss to net photosynthesis,

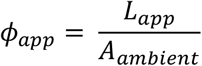

which expresses apparent photorespiratory CO_2_ loss as a fraction of net carbon gain. These quantities are closely related to the difference-based metric used in previous O_2_-shift studies (Fu & Walker, 2024; Sharkey, 1988), but we interpret them explicitly as apparent loss rather than as fully partitioned biochemical rates of photorespiration. Our approach makes several assumptions. First, we assume that mitochondrial day respiration in the light is similar under ambient and N_2_ conditions, such that differences in *A*_net_ primarily reflect changes in photorespiratory flux rather than large changes in respiratory CO_2_ release (Brooks & Farquhar, 1985; Villar *et al*., 1995; Yin & Amthor, 2024). Second, we assume that other sources of CO_2_ release (e.g. from alternative metabolic pathways) are relatively small or change proportionally across our treatments. Third, we assume that short-term exposure to the N_2_ atmosphere does not strongly alter Rubisco activation state (Sage *et al*., 2002) or RuBP regeneration (Schnyder *et al*., 1986) in ways that would dominate the observed differences in *A*_net_.

Under a N_2_ atmosphere, external O_2_ is absent, but the light reactions of photosynthesis still generate O_2_ locally within the leaf through water splitting at PSII under saturating irradiance (Braun, 2020). This internally produced O_2_ allows some Rubisco oxygenation and consequently photorespiration to persist even in nominally anoxic conditions. To minimise exogenous O_2_ availability and thereby suppress photorespiration, a 0% N_2_ environment is preferable to one containing 1–2% O_2_, since higher O_2_ concentrations would increase oxygenation rates and reduce contrast between treatments. However, this approach cannot eliminate photorespiration because internally generated O_2_ sustains residual oxygenation at the site of Rubisco. The occurrence of photorespiration under nominally anoxic conditions is well-documented in photosynthetic microorganisms, where microoxic zones enable O_2_-dependent pathways to function even under overall anoxic conditions (Holert *et al*., 2011; Milrad *et al*., 2023). Two critical limitations therefore apply to our estimates. First, we cannot quantify whether, or to what extent, mitochondrial respiration is suppressed under zero O_2_ conditions *in vivo*; all measurements were made after confirming a new CO_2_ flux steady state on the instrument real-time trace, yet any sustained suppression of mitochondrial respiration under N_2_ would cause ***L***_***app***_ to reflect both reduced photorespiration and changes in respiratory CO_2_ release. Second, because O_2_ is continuously generated internally via the light reactions even under N_2_, our calculated apparent loss likely underestimates total *in vivo* photorespiratory CO_2_ release. Despite these limitations, the O_2_-shift-based difference in *A*_net_ provides a tractable, *in situ* measure of apparent photorespiratory CO_2_ loss across species and temperatures, suitable for assessing relative variation and temperature sensitivity among temperate trees (Tiwari *et al*., 2026).

### ETR partitioning calculations

For each ambient–N_2_ measurement pair, we used the instrument-derived estimates of linear electron transport rate (ETR) to quantify how ambient electron transport was partitioned between net CO_2_ assimilation and apparent photorespiratory CO_2_ loss. Ambient electron transport, *ETR*_ambient_, was taken from chlorophyll fluorescence under 21% O_2_, and ***ETR***_***N***2_, from measurements under the N_2_ atmosphere at otherwise identical conditions. We first calculated the difference in electron transport between ambient and N_2_ conditions as Δ***ETR*** = ***ETR***_***ambient***_ − ***ETR***_***N***2_, which serves as an indicator of O_2_-dependent component of electron use.

To express electron allocation as fractions of ambient ETR, we defined the fraction of ambient electron transport associated with net CO_2_ assimilation (***f***_***Anet***_) and with apparent photorespiratory CO_2_ loss (***f***_***Loss***_) as

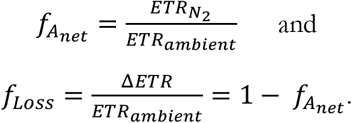

where, ***f***_***Anet***_ represents the fraction of ambient electron transport associated with net CO_2_ assimilation under strongly suppressed photorespiration, and ***f***_***Loss***_ represents the complementary fraction associated with apparent photorespiratory CO_2_ loss and other O_2_-dependent sinks.

We further derived an ETR-based apparent photorespiratory loss term, ***R***_***pETR***_, from the O2-dependent component of electron transport assuming two electrons per CO_2_ released such that

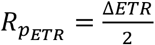

We also calculated the proportional ETR-based apparent loss as

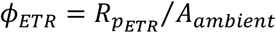

which provides an electron-transport-based analogue of ***ϕ*** = ***L***_***app***_⁄***A***_***ambient***_. These variables were used to assess how the partitioning of electron transport between net assimilation and apparent photorespiratory changed with temperature and among species.

Meaningful comparison of ETR partitioning across species and temperatures requires restricting the analysis to measurement pairs in which the ambient–N_2_ shift can be reasonably interpreted as isolating photorespiratory electron flow, rather than reflecting an artefact of the treatment itself (Gregory et al., 2021; Sharkey, 1985; Sharkey & Vassey, 1989). Under conditions such as saturating light, elevated CO_2_ concentrations, or low temperatures, abrupt exposure to a low-oxygen atmosphere can plausibly drive an immediate cessation of photorespiration, since the oxygenase activity of Rubisco fundamentally requires atmospheric oxygen to function, and its suppression becomes more readily attainable when these conditions favour carboxylation over oxygenation (Migge et al., 1999; Powles & Osmond, 1979). However, this sudden metabolic shift can transiently suppress net photosynthesis by inducing a triose phosphate utilisation (TPU) limitation: the abrupt surge of assimilates overwhelms starch synthesis capacity, trapping inorganic phosphate and temporarily bottlenecking the Calvin-Benson cycle (Cornic & Louason, 1980; McVetty & Canvin, 1981; Rogers et al., 2021; Sharkey & Vassey, 1989).

Most prior reports of this effect used step changes to low, non-zero O_2_ concentrations (e.g., ∼2% O_2_; Sharkey & Vassey, (1989)); our complete removal of O_2_ under N_2_ represents a more extreme version of the same shift and likely increases the probability of TPU limitation rather than reducing it. Hence, in cases where such suppression of both processes occurs, ETR partitioning comparisons are not meaningful for that pair. These specific instances are therefore excluded from our ETR partitioning and temperature-response analyses of *A*_net_ and *L*_app_, which include only measurement pairs where net photosynthesis was not itself confounded by TPU limitation under the N_2_ treatment. (See Supplementary Section N1 for details). The frequency of these exclusions declined with increasing leaf temperature and varied marginally among species (Supplementary Section N1), a pattern consistent with the known temperature sensitivity of TPU limitation (Labate & Leegood, 1988; McClain & Sharkey, 2019; Sage & Sharkey, 1987; Stitt, 1986). ETR partitioning analyses were based on uneven numbers of retained measurement pairs across temperature treatments and species, which should be considered when interpreting comparative patterns.

### Data analysis

To address the first two research questions - examining variation in apparent photorespiratory CO_2_ loss (*L*_app_) across species and temperature, we used mixed-effects models including species, temperature, and their interaction as fixed effects, with individual trees included as a random effect to account for systematic differences among replicates (Supplementary Table S2). Linear models were fitted to visualise temperature responses of both *A*_net_ and ***L***_***app***_. To evaluate whether the O_2_ shift affected stomatal response, we calculated the difference in stomatal conductance between ambient and N_2_ conditions (Δ*g*_s_) for each measurement pair and tested for effects of species and leaf temperature on Δ*g*_s_ using the same mixed-effects model, following the same model structure used for *L*_app_ and *A*_net_ (Supplementary Figure S2). To address the third question on electron transport partitioning, we examined ETR under ambient and N_2_ conditions by analysing ***ETR***_***ambient***_, ***ETR***_***N***2_, and their difference Δ***ETR*** = ***ETR***_***ambient***_ − ***ETR***_***N***2_ for each measurement pair, and related Δ***ETR***, ***f***_***Anet***_, ***f***_***Loss***_, ***R***_***pETR***_, and ***ϕ***_***ETR***_ to temperature using linear regression across all measurement pairs (Supplementary Table S3). All analyses were conducted using R version 4.6.0 (R Core Team, 2026).

## Results

Rates of photosynthesis, *A*_net_, significantly varied across species (*F*_(6, 86)_ = 18.2, *P* <0.001), with values ranging between 5.8 and 17.2 μ mol CO_2_ m^−2^ s^−1^ across the species means and temperature treatments. *B. pendula* had the highest observed mean value for *A*_net_ (17.2 ± 1.3 μ mol CO_2_ m^−2^ s^−1^ at 25 °C) whereas *C. avellana* had the lowest (5.82 ± 0.57 μ mol CO_2_ m^−2^ s^−1^ at 35 °C) (Figure 1, top row; Supplementary Table S4). Across species, the effect of temperature on *A*_net_ was not significant (*F*_(1, 115)_ = 0.01, *P* =0.92), and the Species × *T*_leaf_ interaction was also not significant (*F*_(6, 79)_ = 1.38, *P* =0.23) (Supplementary Table S2).

**Figure 1:**
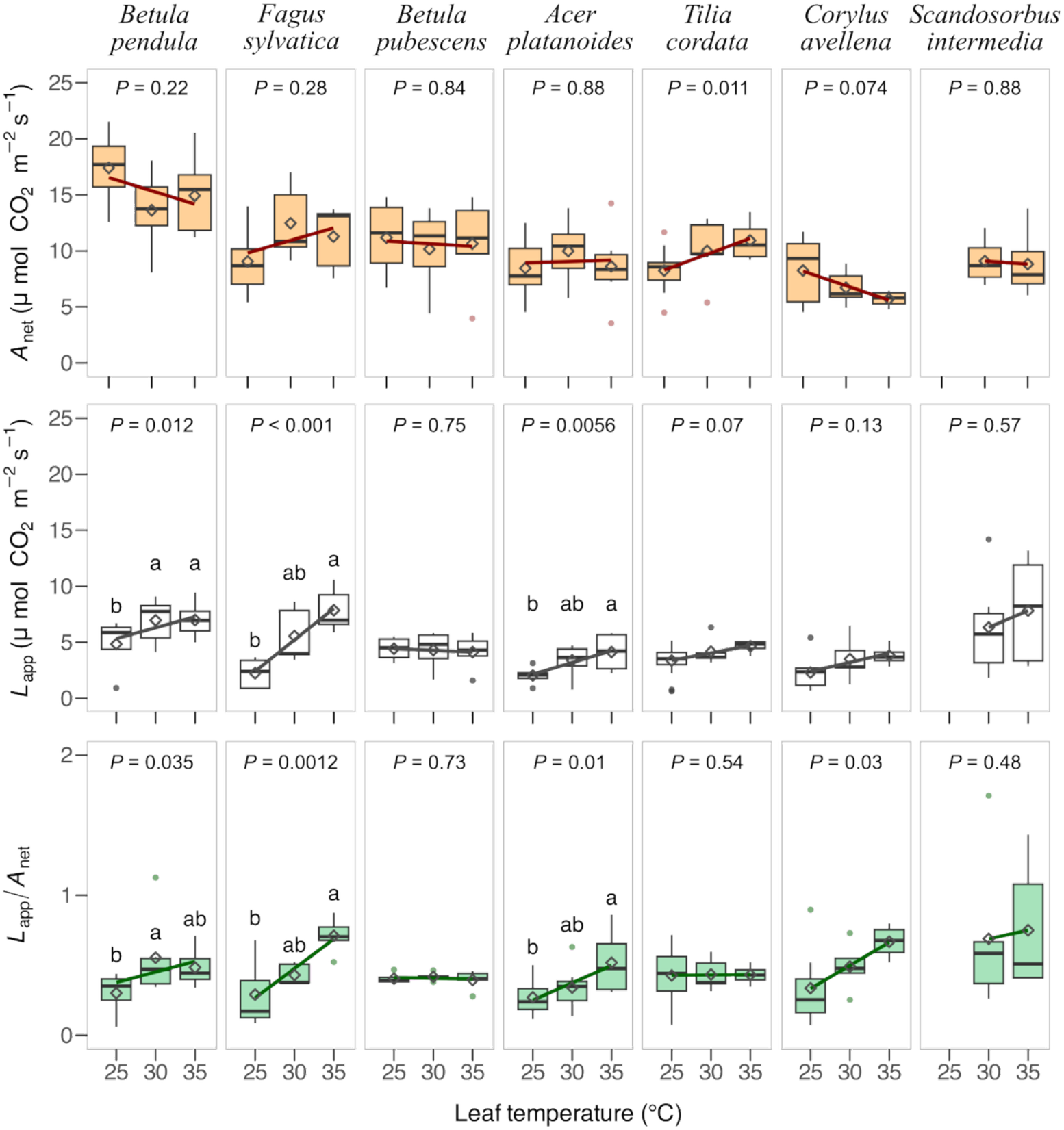
Temperature-dependent variations in photosynthetic CO_2_ assimilation (*A*_net_), apparent photorespiration (*L*_app_) rates, and their ratio (***ϕ*** = *L*_app_/*A*_net_) across seven temperate broadleaf tree species during peak summer conditions measured *in situ* in Uppsala, Sweden. Measurements were taken under saturating irradiance of 1,500 μ mol m^−2^ s^−1^, ambient CO_2_ concentration (420 μ mol mol^−1^) and 50–60% *RH*. *P*-values indicate significance of linear temperature responses – regression lines shown where significant. Note that data for *S. intermedia* at 25°C is missing due to problems during measurement (Supplementary Tables S4 and S5). Letter annotations indicate statistically significant differences between means of temperature points within species. Letters are omitted when there was no significant trend with temperature.

Apparent photorespiratory CO_2_ loss, *L*_app_, showed significant variation among species (*F*_(6, 86)_ = 5.73, *P* <0.001) and temperature treatments (*F*_(1, 115)_ = 26.17, *P* <0.001) (Figure 1, middle row; Supplementary Table S2). Across the three temperature points measured, species-level mean *L*_app_ values ranged from 1.62 to 7.86 μ mol CO_2_ m^−2^ s^−1^. *Fagus sylvatica* had the highest species-mean value for *L*_app_ (7.86 ± 0.88 μ mol CO_2_ m^−2^ s^−1^ at 35 °C), and the lowest value was measured for *Acer platanoides* (1.62 ± 0.49 μ mol CO_2_ m^−2^ s^−1^ at 25 °C) (Supplementary Table S4). The interaction of species and temperature was not significant (*F*_(6, 79)_ = 1.62, *P* = 0.15), indicating that the overall temperature response of *L*_app_ was broadly similar across species.

In general, apparent loss increased with leaf temperature, although the magnitude of the response differed among species. *Fagus sylvatica*, *Betula pendula*, *Acer platanoides*, and *Tilia cordata* showed clear increases in *L*_app_ from 25 to 35 °C, whereas *Betula pubescens* changed little across the temperature treatments. *Corylus avellana* also showed an increasing trend, from 2.31 ± 0.60 to 3.83 ± 0.48 μ mol CO_2_ m^−2^ s^−1^, while *Scandosorbus intermedia* showed high values at 30 and 35 °C but was not measured at 25 °C.

The ratio of apparent loss to net photosynthesis, ***ϕ*** = ***L***_***app***_/***A***_***net***_, also varied significantly across species (*F*_(6, 86)_ = 4.3, *P* <0.001) and increased with temperature (*F*_(1, 115)_ = 17.6, *P* <0.001), whereas the species × *T*_leaf_ interaction was not significant (*F*_(6, 79)_ = 0.9, *P* =0.49) (Figure 1, bottom row; Supplementary Table S2). Across species, mean ***ϕ*** values ranged from 0.18 to 0.41 at 25 °C, 0.34 to 0.69 at 30 °C, and 0.39 to 0.94 at 35 °C. The highest observed value occurred in *Scandosorbus intermedia* at 35 °C (0.94 ± 0.25), followed by *Fagus sylvatica* (0.71 ± 0.06) and *Corylus avellana* (0.67 ± 0.06). These results indicate that the proportional carbon cost of apparent photorespiration increased strongly with temperature.

Stomatal conductance (*g*_s_) showed only modest differences between ambient O_2_ and N_2_ across species and temperatures (Supplementary Figure S2). Across all measurements, Δ*g*_s_ did not differ significantly between species or leaf temperatures (all *P* > 0.2), indicating that the O_2_ shift primarily affected biochemical rather than stomatal processes.

Photosynthetic electron transport also showed strong, condition-dependent responses. Under N_2_, linear electron transport rate was consistently lower than under ambient O_2_ across all species and temperatures, with this reduction becoming progressively stronger at higher leaf temperatures, reflecting the increasing suppression of O_2_-dependent electron sinks under warming. Linear electron transport rate (ETR) differed significantly between ambient and N_2_ conditions and across temperatures (Figure 2). The change in ETR between ambient and N_2_ (ΔETR) varied among species (*F* _(6, 109)_ = 11.3, *P* <0.001) and increased with leaf temperature (*F* _(1, 109)_ = 20.6, *P* <0.001), indicating that suppression of photorespiration under N_2_ had stronger effects on electron transport at warmer temperatures.

**Figure 2:**
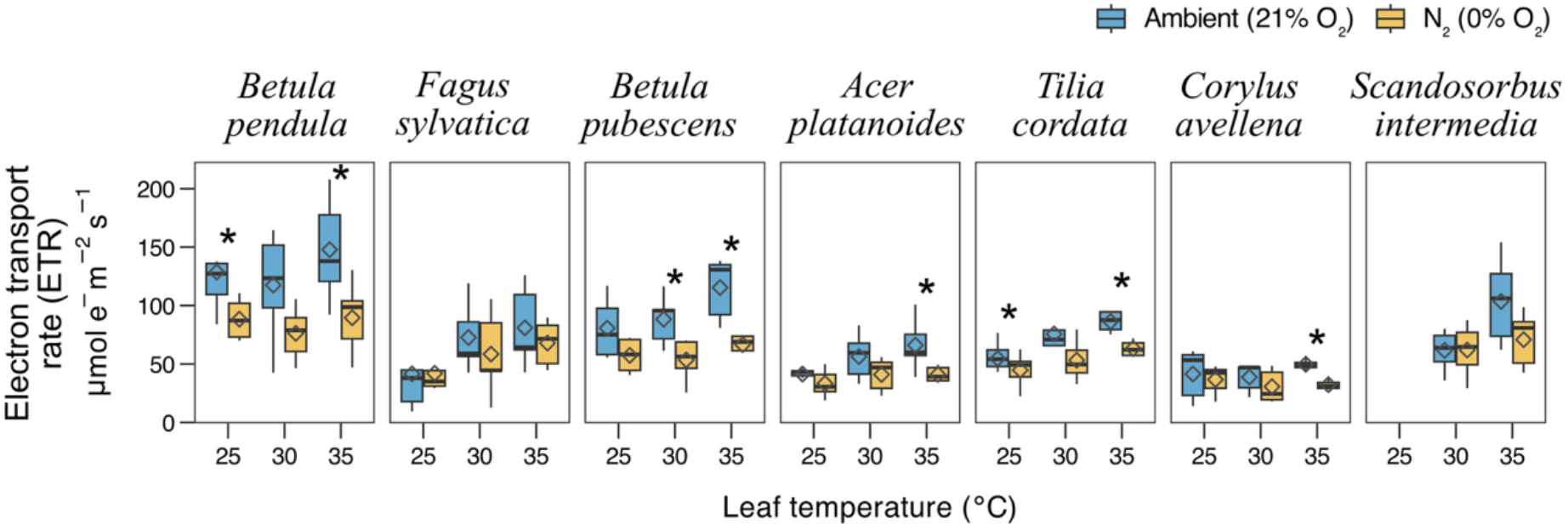
Linear electron transport rate (ETR) responses to ambient and N_2_ conditions across species and temperatures. ETR was measured for seven temperate broadleaf tree species at leaf temperatures of 25, 30, and 35 °C under ambient O_2_ (21% O_2_, blue) and N_2_ (0% O_2_, gold). Diamonds represent mean values. ETR showed pronounced species- and temperature-dependent differences between ambient and N_2_ (ΔETR: species *F* _(6, 109)_ =11.3, *P* <0.001; temperature *F* _(6, 109)_ = 20.6, *P* <0.001), with letter annotations (a, b) indicating significant differences among temperatures within species due to suppression of photorespiration under N_2_.

Partitioning of ambient ETR between net assimilation and apparent loss showed complementary temperature trends (Figure 3). The fraction of ambient electron transport allocated to net CO_2_ assimilation, ***f***_***Anet***_, decreased with temperature (slope = −0.01 °C^−1^, *P* =0.005), while the complementary fraction allocated to apparent photorespiratory CO_2_ loss, ***f***_***Loss***_, increased with equal and opposite slope (≈ +0.01 °C^−1^, *P* =0.005). Across species, ETR partitioning showed a consistent temperature-dependent reallocation of electrons: the fraction of ambient electron transport associated with net CO_2_ assimilation (***f***_***Anet***_) decreased with leaf temperature, while the complementary fraction associated with apparent photorespiratory CO_2_ loss (***f***_***Loss***_), along with the ETR-based apparent loss term (***R***_***p***,***ETR***_) and its proportional expression (***ϕ***_***ETR***_), increased with warming (Supplementary Table S3). ETR-based apparent loss, ***R***_***p***,***ETR***_, rose by ≈ 0.40 μ mol e^−1^ m^−2^ s^−1^ per °C, and the proportional ETR-based apparent loss, ***ϕ***_***ETR***_ = ***R***_***p***,***ETR***_/***A***_***net***,***ambient***_, increased by ≈ 0.04 per °C. Because *f*_Anet_ and *f*_Loss_ are complementary by definition, their opposing trends are not independent evidence; the corroborating result is that the absolute magnitude of ΔETR and the derived ***R***_***p***,***ETR***_ term both increased with temperature, indicating a genuine warming-driven increase in the O_2_-dependent component of electron transport rather than a redistribution artifact of the ratio definition.

**Figure 3:**
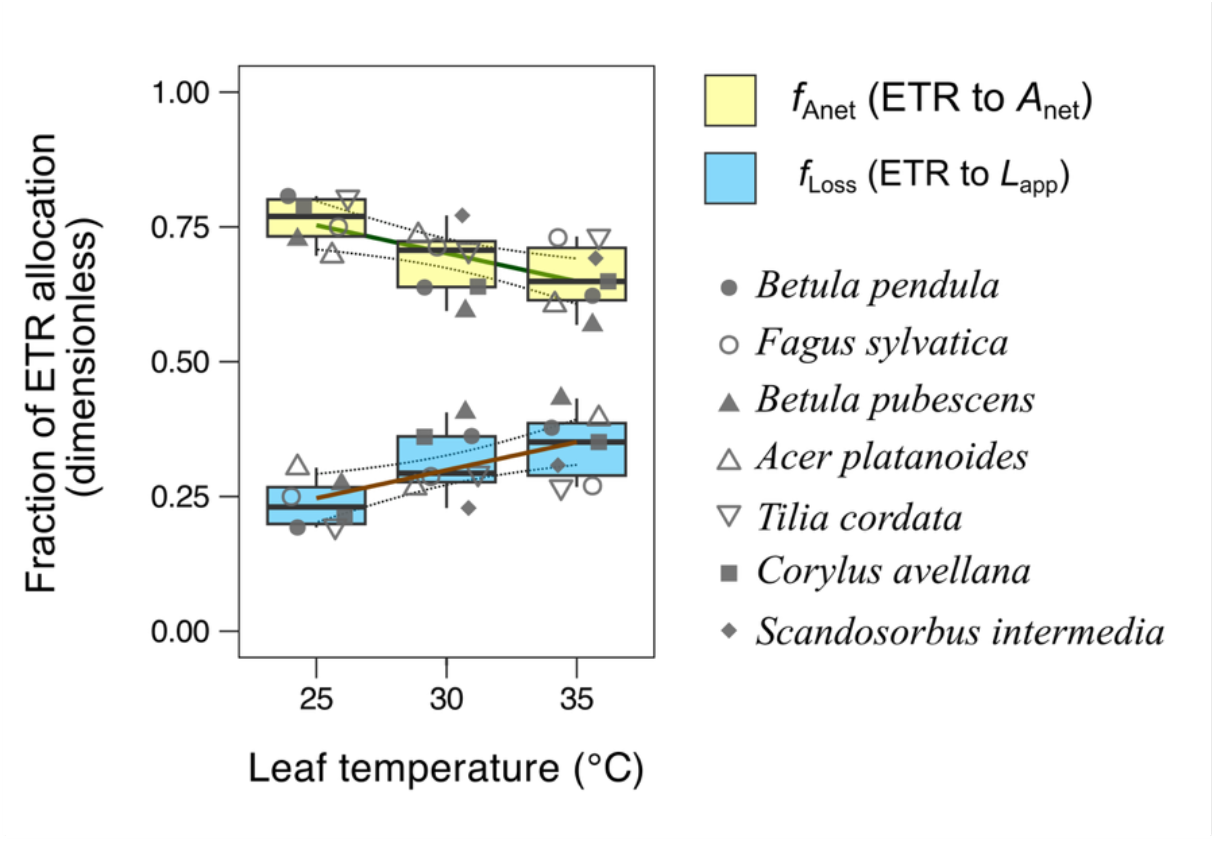
Fraction of linear electron transport rate (ETR) allocated to net CO_2_ assimilation (*A*_net_) and apparent photorespiratory CO_2_ loss (*L*_app_) across temperatures. The panel shows the fraction of ambient electron transport rate (ETR) associated with net CO_2_ assimilation (*f*_Anet_, yellow; boxplots and regression line) and with apparent photorespiratory CO_2_ loss (*f*_Loss_, blue) at leaf temperatures of 25, 30, and 35 °C, calculated from pairs of ambient and N_2_ measurements for seven temperate broadleaf tree species. Points represent species means at each temperature (symbols as indicated in the legend), and boxes show the distribution across measurement pairs. Across species, *f*_Anet_ decreases with temperature while *f*_Loss_ increases, indicating that warming systematically shifts electron transport away from net assimilation and towards apparent photorespiratory CO_2_ loss, with complementary changes in the two fractions (linear regression slopes ≈ −0.01 and +0.01 per °C for *f*_Anet_ and *f*_Loss_, respectively; p ≈ 0.005). Because *f*_Loss_ and *f*_Anet_ are defined as complementary fractions of ambient electron transport, their temperature regressions are mirror images: as leaf temperature increases, the fraction of ETR allocated to net assimilation (*f*_Anet_) decreases while the fraction allocated to apparent photorespiratory CO_2_ loss (*f*_Loss_) increases with equal and opposite slopes and the same statistical strength (|slope| = 0.01 °C^−1^, *P* = 0.005).

The ratio of net CO_2_ assimilation rate to intercellular CO_2_ concentration (*A*_net_/*C*_i_) further illustrated the temperature-dependent impact of photorespiration on assimilation efficiency (Supplementary Figure S3). Species-pooled *A*_net_/*C*_i_ under ambient O_2_ was low and only weakly temperature-dependent, averaging 0.040 at 25 °C, 0.037 at 30 °C, and 0.038 at 35 °C, whereas suppression of photorespiration under N_2_ maintained substantially higher *A*_net_/*C*_i_ at warm temperatures, with means of 0.064, 0.078, and 0.079 at 25, 30, and 35 °C, respectively. At 35 °C, N_2_ *A*_net_/*C*_i_ was on average 0.041 units (≈ 110%) higher than under ambient O_2_, and most species showed their largest ambient–N_2_ separation at this temperature, consistent with stronger photorespiratory limitation of net photosynthesis at high leaf temperatures.

## Discussion

Our key findings include substantial interspecific variation in apparent photorespiratory CO_2_ loss, a general increase in this loss with rising leaf temperature, and a strong temperature-dependent increase in the proportional carbon cost of photorespiration (**ϕ** = *L*_app_/*A*_net_). These patterns are consistent with enhanced Rubisco oxygenation and lower CO_2_ solubility under heat, and they highlight that photorespiration can represent a large fraction of net carbon gain under warm conditions.

### Photorespiration rates vary across temperate trees species

Our results demonstrate previously underrecognized interspecific diversity in photorespiratory metabolism among temperate trees. This variation directly challenges the current practice of parameterizing vegetation models with uniform, crop-derived photorespiration values and suggests that species-specific constraints on Rubisco oxygenation may represent a critical source of functional diversity in forest photosynthetic capacity (Archontoulis et al., 2012; Bernacchi et al., 2001; Han et al., 2020; A. P. Walker et al., 2021).

Photosynthesis and photorespiration rates among the study species were largely consistent with predictions based on the successional associations documented in the literature (Hagemeier & Leuschner, 2019b; Leuschner & Hagemeier, 2020). In particular, rates were mostly higher for acquisitive, early successional species and lower for mid- and late-successional species. However, *F. sylvatica* – a shade-tolerant, late-successional species – exhibited photorespiration rates comparable to the early-successional *B. pendula*, particularly under heat stress. This deviation suggests that photorespiration is influenced by factors beyond successional status and hints at underlying functional diversity in Rubisco properties among species. Interspecific differences in mesophyll conductance or internal CO_2_ supply, which modulate the CO_2_/O_2_ ratio at the Rubisco site, are stronger candidate explanations for the variation observed and remain to be quantified across these species. However, the pronounced species-level divergence in the ratio of net CO_2_ assimilation rate and intercellular CO_2_ concentration (*A*_net_/*C*_i_) under N_2_, - which was not apparent under ambient O_2_, suggests that at least part of the interspecific variation reflects differential biochemical responses to photorespiration suppression rather than differences in *C*_i_ alone (Supplementary Figure S3).

The ratio of apparent photorespiratory CO_2_ loss to net photosynthesis (ϕ = *L*_app_/*A*_net_) provides a proxy metric for integrating photorespiration into physiology models, broadly analogous to the ratio of Rubisco oxygenation and carboxylation (Fernie & Bauwe, 2020; Trudeau et al., 2018). At moderate temperatures (25°C), values ranged from 0.18 to 0.41 (mean 0.3 ± 0.04) across species, somewhat lower than the literature average for woody species (0.56 ± 0.05, Box 1) at 25°C, though rising to 0.48 ± 0.04 at 30°C and 0.59 ± 0.07 at 35°C, approaching the literature baseline as temperatures increase; however, because the literature values are corrected for alternative CO_2_ release pathways while our estimates represent uncorrected apparent loss, this convergence should be interpreted as a qualitative trend rather than a direct quantitative equivalence.

However, species-level variation was substantial, with the lowest values indicating efficient carbon assimilation and the highest reflecting greater proportional investment in photorespiration. Particularly, ϕ increased markedly with temperature, with species mean values reaching up to 0.94 in *S. intermedia* at 35°C - approaching unity and indicating that the apparent carbon cost of photorespiration represented nearly the entirety of net carbon gain under heat stress conditions, even as net CO_2_ assimilation remained positive.

This temperature-driven variation in ϕ has direct implications for FvCB photosynthesis model, foundational to dynamic global vegetation models. The FvCB framework typically assumes fixed photorespiration parameters derived from crop data, yet our results demonstrate this fails to capture interspecific variation and strong temperature sensitivity in temperate trees. Since photorespiration parameters are among the most influential in FvCB predictions, and uncertainty propagates to errors in estimating *V*_cmax_ and *J*_max_, incorporating species-specific and temperature-sensitive ϕ values is essential to improve model fidelity and reduce systematic biases in projecting forest carbon dynamics under warming (Han et al., 2020; Lochocki & McGrath, 2025; B. J. Walker et al., 2017; Z.-P. Ye et al., 2025).

### Temperature sensitivity of photorespiration

Temperate species tend to exhibit photosynthetic responses that remain efficient across a wide range of temperatures, reflecting physiological adaptations to variable thermal environments that allow them to maintain high photosynthetic rates over a broader temperature range (Charles-Edwards & Charles-Edwards, 1970; Crous *et al*., 2022). In our study, *A*_net_ typically declined slightly or remained unchanged with increasing temperature, which may indicate that the thermal optimum was either already exceeded at 25°C or not reached at 35°C. With only three temperature points, it is not possible to precisely determine the thermal optimum. However, although not statistically significant, three species – *F. sylvatica*, *A. platanoides*, and *T. cordata* – displayed higher *A*_net_ rates at 30 °C – than at 25 °C or 35 °C. Nonetheless, leaves generally maintained high *A*_net_ rates even at 35 °C. Further experiments are required to clarify the shape of these temperature response curves.

In contrast to the largely invariant temperature response of *A*_net_ across species, apparent photorespiratory CO_2_ loss (*L*_app_) increased with temperature in four species, while in three species it did not change. In *C. avellena*, the trend of increasing *L*_app_ with temperature was apparent but not statistically significant. Thus, some species showed a clear temperature-driven increase in photorespiration, consistent with a protective role under thermal stress conditions (Brooks & Farquhar, 1985; Ciereszko & Kuźniak, 2024; Jordan & Ogren, 1984; Morales et al., 2020), whereas in others the increase was weak or absent, and in *B. pubescens* there was no temperature effect on photorespiration. The physiological significance of this insensitivity remains unclear, but it may reflect interspecific differences in thermal stress responses, in the contribution of photorespiration to protection, or in the use of alternative protective mechanisms that are not yet understood.

In our study, the average ratio of apparent photorespiratory CO_2_ release to net photosynthesis (*L*_app_/*A*_net_) increased from 0.18 at 25 °C to an average of 0.59, in some cases going up to 0.93 at 35 °C. *In vitro* enzyme kinetic studies show that the oxygenation to carboxylation ratio (*V*_O_/*V*_C_) of Rubisco in wheat rises from 0.25 at 10 °C to 0.56 at 35 °C, under conditions simulating sub-atmospheric CO_2_ concentrations (Brooks & Farquhar, 1985; Hall & Keys, 1983). If the enzymatic Rubisco oxygenation rate reflects leaf-level photorespiratory flux, the temperature sensitivity (slope) of *V*_O_/*V*_C_ is 0.0124 °C^−1^ (from 0.25 to 0.56 across 10–35 °C), while the increase in *R*_p_/*A*_net_ observed in our measurements is steeper, at 0.038 °C⁻¹ (from 0.52 to 0.93 between 25–35 °C). This suggests that the leaf-level photorespiratory response to temperature may be more sensitive than indicated by isolated Rubisco enzyme kinetics. However, we note that this comparison assumes a close correspondence between enzyme kinetics and leaf-level fluxes, which remains to be tested in forest trees. While the slope of the *L*_app_/*A*_net_ varied among species studied, distinctly, the late-successional species *F. sylvatica* showed a pronounced increase in *L*_app_ at elevated temperatures, exceeding that of mid-successional species such as *T. cordata* and *A. platanoides*. Also, *B. pubescens* exhibited no measurable increase in apparent *R*_p_ with temperature.

Considering the role of photorespiration in thermal protection, fast-growing, acquisitive species appear to prioritize carbon acquisition rather than heat defence, whereas slow-growing, late-successional species must prioritise photorespiration and thermal tolerance to safeguard longevity (Bazzaz, 1979; Mujawamariya et al., 2023; Slot & Winter, 2017). Our measured photorespiration rates generally support this trade-off, with the notable exceptions of *F. sylvatica* (a late-successional species with unexpectedly high thermal response) and *B. pubescens* (which showed no temperature effect). The modest changes in stomatal conductance and the consistent decline in *A*_net_/*C*_i_ under ambient O_2_ but not under N_2_ (Supplementary Figures S2 and S3) support the view that warming-induced changes in net photosynthesis are driven primarily by biochemical photorespiratory processes rather than stomatal closure. Whether these divergent responses reflect differing degrees of thermal stress tolerance or genetic adaptation of Rubisco remains to be determined (Hall & Keys, 1983; Kozaki & Takeba, 1996). Incorporating species-specific and temperature-sensitive photorespiration parameters will help refine predictions of forest carbon dynamics (A. Cavanagh & Matthews, 2025; Lochocki & McGrath, 2025; van Bodegom et al., 2014).

### Electron transport partitioning supports a photoprotective role

Our *in situ* O_2_-shift measurements and ETR partitioning provide one of the first field-based, multi-species datasets addressing the knowledge gaps highlighted by Tiwari et al., (2026), particularly the lack of tree-specific constraints on photorespiratory temperature sensitivity and electron transport partitioning. By showing that *ϕ* = *L*_app_/*A*_net_ increases steeply with leaf temperature and that this trend is independently corroborated by a rise in ETR-based apparent loss (***R***_***p***,***ETR***_) and its proportional expression (ϕ*ETR*) – both derived from the absolute magnitude of ΔETR rather than from the *f*_Anet_/*f*_Loss_ partitioning ratio; our results empirically support the view that photorespiration acts as a temperature-sensitive energy sink in temperate trees rather than a purely wasteful flux. This temperature-sensitive energy-sink role of photorespiration is likely especially consequential in northern temperate regions, where short growing seasons and long daylight hours expose trees to extended heat and irradiance, making photoprotection and thermal regulation critical for sustained carbon gain as extreme heat events become more frequent (Perkins-Kirkpatrick & Lewis, 2020; B. J. Walker et al., 2016). Building on the interspecific trade-offs and temperature sensitivity identified above, we selected seven widely distributed broadleaf tree species from the northern temperate region and conducted *in situ* measurements of *R*_p_ and *A*_net_ at three leaf temperatures to test how photorespiration scales with warming.

### Proportional carbon cost of photorespiration increases with warming

Across all species, the ratio of apparent photorespiratory CO_2_ loss to net photosynthesis (***ϕ*** = ***L***_***app***_/***A***_***net***_) increased consistently with leaf temperature (*F*_(1,115)_ = 17.6, *P* < 0.001), indicating that warming progressively elevated the proportional carbon cost of photorespiration relative to net carbon gain. At 35°C, ***ϕ*** reached 0.94 in *S. intermedia*, meaning apparent photorespiratory loss represented nearly the entirety of net carbon gain under heat stress conditions, even as *A*_net_ remained positive in all species and temperatures measured. This sustained net assimilation under high ***ϕ*** values are consistent with a functional role for photorespiration as an energy dissipation pathway under thermal stress: absorbing excess reductant and preventing photodamage when carboxylation is limited (A. P. Cavanagh et al., 2022; Huang et al., 2019; Kozaki & Takeba, 1996; Streb et al., 2005; Wingler et al., 2000). The parallel, independently derived increase in ETR-based apparent loss (***R***_***p***,***ETR***_) with warming (Supplementary Table S3) further supports the interpretation that photorespiration functions as an increasingly important alternative electron sink as leaf temperatures rise toward and beyond thermal optima.

Isotopic fractionation approaches can, in principle, estimate photorespiration rates by partitioning carbon isotope discrimination among carboxylation, photorespiration and day respiration (Busch, 2013; Lloyd et al., 2023; Ubierna & Farquhar, 2014), but have so far been developed and tested mainly in herbaceous C3 species. Our *in situ* O_2_-shift measurements of apparent photorespiratory CO_2_ loss and its temperature sensitivity in temperate broadleaf trees therefore provide complementary leaf-level flux constraints that could help refine future isotope-based applications in forest systems (Ehlers *et al*., 2015).

### Conclusion

We studied leaf-level photorespiration rates across seven temperate broadleaf tree species, measured *in situ* under ecologically relevant temperatures. The results show substantial species wide variation in photorespiration and a generally strong positive response of photorespiration to increasing leaf temperature, along with changes in the ratio of photorespiration to net photosynthesis. Although species displayed diverse thermal responses while maintaining stable photosynthesis, most showed increased photorespiration at higher temperatures, with some exceptions where photorespiration did not increase. The observed increase in the proportional carbon cost of photorespiration and the temperature-dependent reallocation of electron transport away from net CO_2_ assimilation and towards apparent photorespiratory loss under heat stress highlight complex regulatory and protective processes. These findings are important for vegetation models that largely rely on crop-based photorespiration parameters, which tend to underestimate the variation and temperature sensitivity seen in forest species.

## Supporting information

Supplementary Section 3

## Acknowledgement

Research was funded by grants from the Wenner-Gren Foundation (to RT and RM), the Birgitta Sintring Foundation (to RT and RM), the Swedish Phytogeographical Society (to RT and RM), the J.W Palmstruch Scholarship fund (to PD), and the Swedish Research Council, Formas (2020-00921 to RM). We thank Christoffer Bergvall, Adriana Puentes, Gustaf Granath for valuable assistance. We thank H S Shankaranarayana Bhatta, Somashekhara Achar KG for the help with Sanskrit and Kannada language version of the article. We thank Shankaranarayana Bhatta HS, Vasudeva HR, Chethan TR and Pragya Rakesh Tiwari for their help with the Sanskrit, Kannada and Hindi language versions of the manuscript. RT thanks Shāradā-Chandramaulishwara and Ubhaya Jagadgurus of the Dakshinamnāya Sri Sharadā Peetham, Sringēri for their guidance.

## Ethics approval and statements

Not applicable.

## Data availability

Data associated with the manuscript is available on Figshare: 10.6084/m9.figshare.33190686

## Author contributions

RT conceived the study, developed the methodology, carried out the analysis and led the writing of the original draft. PD and RT collected the data. RM supervised the study. All authors contributed to the writing and revision of the manuscript.

## Disclosing artificial intelligence use

Perplexity AI (Perplexity AI, Inc.) was used solely to improve the clarity and language of the manuscript text during drafting; it was not used to generate, analyze, or interpret any data, results, or scientific content.

## References

Archontoulis, S. V., Yin, X., Vos, J., Danalatos, N. G., & Struik, P. C. (2012). Leaf photosynthesis and respiration of three bioenergy crops in relation to temperature and leaf nitrogen: how conserved are biochemical model parameters among crop species? Journal of Experimental Botany, 63(2), 895– 911. 10.1093/jxb/err321

Bauwe, H., Hagemann, M., & Fernie, A. R. (2010). Photorespiration: players, partners and origin. Trends in Plant Science, 15(6), 330–336. 10.1016/j.tplants.2010.03.006

Bazzaz, F. A. (1979). The physiological ecology of plant succession. Annual Review of Ecology and Systematics, 10(1), 351–371. 10.1146/annurev.es.10.110179.002031

Beauclaire, Q., Vanden Brande, F., & Longdoz, B. (2024). Key role played by mesophyll conductance in limiting carbon assimilation and transpiration of potato under soil water stress. Frontiers in Plant Science, 15, 1500624. 10.3389/fpls.2024.1500624

Bernacchi, C. J., Singsaas, E. L., Pimentel, C., Portis, A. R., Jr, & Long, S. P. (2001). Improved temperature response functions for models of Rubisco-limited photosynthesis: *In vivo*Rubisco enzyme kinetics. Plant, Cell & Environment, 24(2), 253–259. 10.1111/j.1365-3040.2001.00668.x

Braun, H.-P. (2020). The Oxidative Phosphorylation system of the mitochondria in plants. Mitochondrion, 53, 66–75. 10.1016/j.mito.2020.04.007

Brooks, A., & Farquhar, G. D. (1985). Effect of temperature on the CO_2_/O _2_ specificity of ribulose-1,5-bisphosphate carboxylase/oxygenase and the rate of respiration in the light: Estimates from gas-exchange measurements on spinach: Estimates from gas-exchange measurements on spinach. Planta, 165(3), 397–406. 10.1007/BF00392238

Busch, F. A. (2013). Current methods for estimating the rate of photorespiration in leaves. *Plant Biology (Stuttgart*, Germany), 15(4), 648–655. 10.1111/j.1438-8677.2012.00694.x

Busch, F. A., Deans, R. M., & Holloway-Phillips, M.-M. (2017). Estimation of photorespiratory fluxes by gas exchange. *Methods in Molecular Biology (Clifton*, N.J.), 1653, 1–15. 10.1007/978-1-4939-7225-8_1

Busch, F. A., & Sage, R. F. (2017). The sensitivity of photosynthesis to O_2_ and CO_2_ concentration identifies strong Rubisco control above the thermal optimum. The New Phytologist, 213(3), 1036–1051. 10.1111/nph.14258

Cavanagh, A., & Matthews, M. (2025). The heat is on: scaling improvements in photosynthetic thermal tolerance from the leaf to canopy to predict crop yields in a changing climate. Philosophical Transactions of the Royal Society of London. Series B, Biological Sciences, 380(1927), 20240235. 10.1098/rstb.2024.0235

Cavanagh, A. P., South, P. F., Bernacchi, C. J., & Ort, D. R. (2022). Alternative pathway to photorespiration protects growth and productivity at elevated temperatures in a model crop. Plant Biotechnology Journal, 20(4), 711–721. 10.1111/pbi.13750

Charles-Edwards, D. A., & Charles-Edwards, J. (1970). An analysis of the temperature response curves of CO_2_ exchange in the leaves of two temperate and one tropical grass species. Planta, 94(2), 140–151. 10.1007/BF00387758

Chuang, J., & Ling, L. (2005). The co-operation of leaf orientation, photorespiration and thermal dissipation alleviate photoinhibition in young leaves of soybean plants. https://www.semanticscholar.org/paper/The-co-operation-of-leaf-orientation%2C-and-thermal-Chuang-Ling/4b873939280af7a6aa5d4b1cec6e905f3b2f0ac4

Ciereszko, I., & Kuźniak, E. (2024). Photorespiratory metabolism and its regulatory links to plant defence against pathogens. International Journal of Molecular Sciences, 25(22), 12134. 10.3390/ijms252212134

Cornic, G., & Louason, G. (1980). The effects of O_2_ on net photosynthesis at low temperature (5°C). Plant, Cell & Environment, 3(2), 149–157. 10.1111/1365-3040.ep11580938

Crous, K. Y., Uddling, J., & De Kauwe, M. G. (2022). Temperature responses of photosynthesis and respiration in evergreen trees from boreal to tropical latitudes. The New Phytologist, 234(2), 353–374. 10.1111/nph.17951

de Rigo, D., Durrant, T., Caudullo, G., & Barredo, J. I. (2016). European forests: an ecological overview. European Atlas of Forest Tree Species, 24–31.

Desotgiu, R., Cascio, C., Pollastrini, M., Gerosa, G., Marzuoli, R., & Bussotti, F. (2012). Short and long term photosynthetic adjustments in sun and shade leaves ofFagus sylvaticaL., investigated by fluorescence transient (FT) analysis. Plant Biosystems, 146(sup1), 206–216. 10.1080/11263504.2012.705350

Diao, H., Cernusak, L. A., Saurer, M., Gessler, A., Siegwolf, R. T. W., & Lehmann, M. M. (2024). Uncoupling of stomatal conductance and photosynthesis at high temperatures: mechanistic insights from online stable isotope techniques. The New Phytologist, 241(6), 2366–2378. 10.1111/nph.19558

Ehlers, I., Augusti, A., Betson, T. R., Nilsson, M. B., Marshall, J. D., & Schleucher, J. (2015). Detecting long-term metabolic shifts using isotopomers: CO_2_-driven suppression of photorespiration in C3 plants over the 20^th^ century. Proceedings of the National Academy of Sciences of the United States of America, 112(51), 15585–15590. 10.1073/pnas.1504493112

Epron, D., Godard, D., Cornic, G., & Genty, B. (1995). Limitation of net CO_2_ assimilation rate by internal resistances to CO_2_ transfer in the leaves of two tree species (*Fagus sylvatica* L. and *Castanea sativa* Mill.). Plant, Cell & Environment, 18(1), 43–51. 10.1111/j.1365-3040.1995.tb00542.x

Farquhar, G. D., von Caemmerer, S., & Berry, J. A. (1980). A biochemical model of photosynthetic CO_2_ assimilation in leaves of C 3 species. Planta, 149(1), 78–90. 10.1007/BF00386231

Fernie, A. R., & Bauwe, H. (2020). Wasteful, essential, evolutionary stepping stone? The multiple personalities of the photorespiratory pathway. The Plant Journal: For Cell and Molecular Biology, 102(4), 666–677. 10.1111/tpj.14669

Foyer, C. H., Bloom, A. J., Queval, G., & Noctor, G. (2009). Photorespiratory metabolism: genes, mutants, energetics, and redox signaling. Annual Review of Plant Biology, 60(1), 455–484. 10.1146/annurev.arplant.043008.091948

Foyer, C. H., Lelandais, M., & Kunert, K. J. (1994). Photooxidative stress in plants. Physiologia Plantarum, 92(4), 696–717. 10.1111/j.1399-3054.1994.tb03042.x

Fu, X., & Walker, B. J. (2024). Using dynamic gas exchange measurements during oxygen transients to study nonsteady-state photorespiration. *Methods in Molecular Biology (Clifton*, N.J.), 2792, 175–184. 10.1007/978-1-0716-3802-6_14

Gregory, L. M., McClain, A. M., Kramer, D. M., Pardo, J. D., Smith, K. E., Tessmer, O. L., Walker, B. J., Ziccardi, L. G., & Sharkey, T. D. (2021). The triose phosphate utilization limitation of photosynthetic rate: Out of global models but important for leaf models. Plant, Cell & Environment, 44(10), 3223–3226. 10.1111/pce.14153

Hagemeier, M., & Leuschner, C. (2019a). Functional crown architecture of five temperate broadleaf tree species: Vertical gradients in leaf morphology, leaf angle, and leaf area density. Forests, 10(3), 265. 10.3390/f10030265

Hagemeier, M., & Leuschner, C. (2019b). Leaf and crown optical properties of five early-, mid- and late-successional temperate tree species and their relation to sapling light demand. Forests, 10(10), 925. 10.3390/f10100925

Hall, N. P., & Keys, A. J. (1983). Temperature dependence of the enzymic carboxylation and oxygenation of ribulose 1,5-bisphosphate in relation to effects of temperature on photosynthesis. Plant Physiology, 72(4), 945–948. 10.1104/pp.72.4.945

Han, T., Zhu, G., Ma, J., Wang, S., Zhang, K., Liu, X., Ma, T., Shang, S., & Huang, C. (2020). Sensitivity analysis and estimation using a hierarchical Bayesian method for the parameters of the FvCB biochemical photosynthetic model. Photosynthesis Research, 143(1), 45–66. 10.1007/s11120-019-00684-z

Hochberg, U., Degu, A., Fait, A., & Rachmilevitch, S. (2013). Near isohydric grapevine cultivar displays higher photosynthetic efficiency and photorespiration rates under drought stress as compared with near anisohydric grapevine cultivar. Physiologia Plantarum, 147(4), 443–452. 10.1111/j.1399-3054.2012.01671.x

Holert, J., Hahnke, S., & Cypionka, H. (2011). Influence of light and anoxia on chemiosmotic energy conservation in *Dinoroseobacter shibae*: Energy conservation in *Dinoroseobacter shibae*. Environmental Microbiology Reports, 3(1), 136–141. 10.1111/j.1758-2229.2010.00199.x

Hu, S., Ding, Y., & Zhu, C. (2020). Sensitivity and responses of chloroplasts to heat stress in plants. Frontiers in Plant Science, 11, 375. 10.3389/fpls.2020.00375

Huang, W., Hu, H., & Zhang, S.-B. (2015). Photorespiration plays an important role in the regulation of photosynthetic electron flow under fluctuating light in tobacco plants grown under full sunlight. Frontiers in Plant Science, 6, 621. 10.3389/fpls.2015.00621

Huang, W., Yang, Y.-J., Wang, J.-H., & Hu, H. (2019). Photorespiration is the major alternative electron sink under high light in alpine evergreen sclerophyllous *Rhododendron* species. Plant Science: An International Journal of Experimental Plant Biology, 289(110275), 110275. 10.1016/j.plantsci.2019.110275

Jiang, X., Walker, B. J., He, S. Y., & Hu, J. (2023). The role of photorespiration in plant immunity. Frontiers in Plant Science, 14, 1125945. 10.3389/fpls.2023.1125945

Jordan, D. B., & Ogren, W. L. (1984). The CO_2_/O _2_ specificity of ribulose 1,5-bisphosphate carboxylase/oxygenase: Dependence on ribulosebisphosphate concentration, pH and temperature. Planta, 161(4), 308–313. 10.1007/BF00398720

Keys, A. J., Sampaio, E. V. S. B., Cornelius, M. J., & Bird, I. F. (1977). Effect of temperature on photosynthesis and photorespiration of wheat leaves. Journal of Experimental Botany, 28(3), 525–533. 10.1093/jxb/28.3.525

Kozaki, A., & Takeba, G. (1996). Photorespiration protects C3 plants from photooxidation. Nature, 384(6609), 557–560. 10.1038/384557a0

Kuppusamy, A., Alagarswamy, S., Karuppusami, K. M., Maduraimuthu, D., Natesan, S., Ramalingam, K., Muniyappan, U., Subramanian, M., & Kanagarajan, S. (2023). Melatonin enhances the photosynthesis and antioxidant enzyme activities of mung bean under drought and high-temperature stress conditions. Plants, 12(13), 2535. 10.3390/plants12132535

Labate, C. A., & Leegood, R. C. (1988). Limitation of photosynthesis by changes in temperature: Factors affecting the response of carbon-dioxide assimilation to temperature in barley leaves: Factors affecting the response of carbon-dioxide assimilation to temperature in barley leaves. Planta, 173(4), 519–527. 10.1007/BF00958965

Leuschner, C., & Hagemeier, M. (2020). The economy of canopy space occupation and shade production in early-to late-successional temperate tree species and their relation to productivity. Forests, 11(3), 317. 10.3390/f11030317

Lloyd, M. K., Stein, R. A., Ibarra, D. E., Barclay, R. S., Wing, S. L., Stahle, D. W., Dawson, T. E., & Stolper, D. A. (2023). Isotopic clumping in wood as a proxy for photorespiration in trees. Proceedings of the National Academy of Sciences of the United States of America, 120(46), e2306736120. 10.1073/pnas.2306736120

Lochocki, E. B., & McGrath, J. M. (2025). Widely used variants of the Farquhar-von-Caemmerer-Berry model can cause errors in parameter estimation. In Silico Plants, 7(diaf014), diaf014. 10.1093/insilicoplants/diaf014

Lorimer, G. H., & Andrews, T. J. (1973). Plant photorespiration—an inevitable consequence of the existence of atmospheric oxygen. Nature, 243(5406), 359–360. 10.1038/243359a0

Marchin, R. M., Medlyn, B. E., Tjoelker, M. G., & Ellsworth, D. S. (2023). Decoupling between stomatal conductance and photosynthesis occurs under extreme heat in broadleaf tree species regardless of water access. Global Change Biology, 29(22), 6319–6335. 10.1111/gcb.16929

McClain, A. M., & Sharkey, T. D. (2019). Triose phosphate utilization and beyond: from photosynthesis to end product synthesis. Journal of Experimental Botany, 70(6), 1755–1766. 10.1093/jxb/erz058

McVetty, P. B. E., & Canvin, D. T. (1981). Inhibition of photosynthesis by low oxygen concentrations. Canadian Journal of Botany, 59(5), 721–725. 10.1139/b81-102

Migge, A., Kahmann, U., Fock, H. P., & Becker, T. W. (1999). Prolonged exposure of tobacco to a low oxygen atmosphere to suppress photorespiration decreases net photosynthesis and results in changes in plant morphology and chloroplast structure. Photosynthetica, 36(1–2), 107–116. 10.1023/a:1007074921833

Milrad, Y., Nagy, V., Elman, T., Fadeeva, M., Tóth, S. Z., & Yacoby, I. (2023). A PSII photosynthetic control is activated in anoxic cultures of green algae following illumination. Communications Biology, 6(1), 514. 10.1038/s42003-023-04890-3

Morales, F., Ancín, M., Fakhet, D., González-Torralba, J., Gámez, A. L., Seminario, A., Soba, D., Ben Mariem, S., Garriga, M., & Aranjuelo, I. (2020). Photosynthetic metabolism under stressful growth conditions as a bases for crop breeding and yield improvement. Plants, 9(1), 88. 10.3390/plants9010088

Mujawamariya, M., Wittemann, M., Dusenge, M. E., Manishimwe, A., Ntirugulirwa, B., Zibera, E., Nsabimana, D., Wallin, G., & Uddling, J. (2023). Contrasting warming responses of photosynthesis in early- and late-successional tropical trees. Tree Physiology, 43(7), 1104–1117. 10.1093/treephys/tpad035

Noctor, G., Veljovic-Jovanovic, S., Driscoll, S., Novitskaya, L., & Foyer, C. H. (2002). Drought and oxidative load in the leaves of C3 plants: a predominant role for photorespiration? Annals of Botany, 89 Spec No(SI), 841–850. 10.1093/aob/mcf096

Oberpriller, J., Herschlein, C., Anthoni, P., Arneth, A., Krause, A., Rammig, A., Lindeskog, M., Olin, S., & Hartig, F. (2022). Climate and parameter sensitivity and induced uncertainties in carbon stock projections for European forests (using LPJ-GUESS 4.0). Geoscientific Model Development, 15(16), 6495–6519. 10.5194/gmd-15-6495-2022

Ogren, W. L., & Bowes, G. (1971). Ribulose diphosphate carboxylase regulates soybean photorespiration. Nature: New Biology, 230(13), 159–160. 10.1038/newbio230159a0

Osei-Bonsu, I., McClain, A. M., Walker, B. J., Sharkey, T. D., & Kramer, D. M. (2021). The roles of photorespiration and alternative electron acceptors in the responses of photosynthesis to elevated temperatures in cowpea. Plant, Cell & Environment, 44(7), 2290–2307. 10.1111/pce.14026

Õunapuu-Pikas, E., Tullus, A., Kupper, P., Tamm, I., Reinthal, T., & Sellin, A. (2025). Foliage development and resource allocation determine the growth responses of silver birch (*Betula pendula*) to elevated environmental humidity. Tree Physiology, 45(1). 10.1093/treephys/tpae161

Perkins-Kirkpatrick, S. E., & Lewis, S. C. (2020). Increasing trends in regional heatwaves. Nature Communications, 11(1), 3357. 10.1038/s41467-020-16970-7

Peterson, R. B. (1989). Partitioning of noncyclic photosynthetic electron transport to O(_2)_-dependent dissipative processes as probed by fluorescence and CO(_2)_ exchange. Plant Physiology, 90(4), 1322– 1328. 10.1104/pp.90.4.1322

Powles, S. B., & Osmond, C. B. (1979). Photoinhibition of intact attached leaves of c(3) plants illuminated in the absence of both carbon dioxide and of photorespiration. Plant Physiology, 64(6), 982–988. 10.1104/pp.64.6.982

R Core Team,. (2026). R: A language and environment for statistical computing (Version 4.6.0). https://www.R-project.org/

Rogers, A., Kumarathunge, D. P., Lombardozzi, D. L., Medlyn, B. E., Serbin, S. P., & Walker, A. P. (2021). Triose phosphate utilization limitation: an unnecessary complexity in terrestrial biosphere model representation of photosynthesis. The New Phytologist, 230(1), 17–22. 10.1111/nph.17092

Rokka, A., Aro, E. M., Herrmann, R. G., Andersson, B., & Vener, A. V. (2000). Dephosphorylation of photosystem II reaction center proteins in plant photosynthetic membranes as an immediate response to abrupt elevation of temperature. Plant Physiology, 123(4), 1525–1536. 10.1104/pp.123.4.1525

Roze, L. V., Antoniak, A., Sarkar, D., Liepman, A. H., Tejera-Nieves, M., Vermaas, J. V., & Walker, B. J. (2025). Increasing thermostability of the key photorespiratory enzyme glycerate 3-kinase by structure-based recombination. Plant Biotechnology Journal, 23(2), 454–466. 10.1111/pbi.14508

Rydin, H., Snoeijs, P., & Diekmann, M. (1999). *Swedish plant geography* (H. Rydin, P. Snoeijs, & M. Diekmann, Eds). Svenska Vaxtgeografiska Sallskapet.

Sage, R. F., Cen, Y.-P., & Li, M. (2002). The activation state of Rubisco directly limits photosynthesis at low CO(2) and low O(2) partial pressures. Photosynthesis Research, 71(3), 241–250. 10.1023/A:1015510005536

Sage, R. F., & Sharkey, T. D. (1987). The effect of temperature on the occurrence of O(2) and CO(2) insensitive photosynthesis in field grown plants. Plant Physiology, 84(3), 658–664. 10.1104/pp.84.3.658

Salvucci, M. E., & Crafts-Brandner, S. J. (2004). Inhibition of photosynthesis by heat stress: the activation state of Rubisco as a limiting factor in photosynthesis. Physiologia Plantarum, 120(2), 179–186. 10.1111/j.0031-9317.2004.0173.x

Schnyder, H., Mächler, F., & Nösberger, J. (1986). Regeneration of Ribulose 1,5-bisphosphate and Ribulose 1,5-bisphosphate carboxylase/oxygenase Activity Associated with Lack of Oxygen Inhibition of Photosynthesis at Low Temperature. Journal of Experimental Botany, 37(8), 1170–1179. 10.1093/jxb/37.8.1170

Sharkey, T. D. (1985). O2-Insensitive Photosynthesis in C3 Plants. Plant Physiology, 78(1), 71–75. https://www.ncbi.nlm.nih.gov/pubmed/16664211

Sharkey, T. D. (1988). Estimating the rate of photorespiration in leaves. Physiologia Plantarum, 73(1), 147–152. 10.1111/j.1399-3054.1988.tb09205.x

Sharkey, T. D., Berry, J. A., & Sage, R. F. (1988). Regulation of photosynthetic electron-transport in *Phaseolus vulgaris* L., as determined by room-temperature chlorophyll a fluorescence. Planta, 176(3), 415–424. 10.1007/BF00395423

Sharkey, T. D., & Vassey, T. L. (1989). Low oxygen inhibition of photosynthesis is caused by inhibition of starch synthesis. Plant Physiology, 90(2), 385–387. 10.1104/pp.90.2.385

Shi, Q., Sun, H., Timm, S., Zhang, S., & Huang, W. (2022). Photorespiration alleviates photoinhibition of photosystem I under fluctuating light in tomato. Plants, 11(2), 195. 10.3390/plants11020195

Slot, M., & Winter, K. (2017). Photosynthetic acclimation to warming in tropical forest tree seedlings. Journal of Experimental Botany, 68(9), 2275–2284. 10.1093/jxb/erx071

SMHI. (2024). Meteorological Observations from Station 97510 (Uppsala Aut) [Data set]. In Swedish Meteorological and Hydrological Institute (SMHI). https://www.smhi.se/data/meteorologi/ladda-ner-meteorologiska-observationer#param=airHumidity,stations=core,stationid=97510

Stitt, M. (1986). Limitation of photosynthesis by carbon metabolism: I. evidence for excess electron transport capacity in leaves carrying out photosynthesis in saturating light and CO(2): I. Evidence for excess electron transport capacity in leaves carrying out photosynthesis in saturating light and CO2. Plant Physiology, 81(4), 1115–1122. 10.1104/pp.81.4.1115

Streb, P., Josse, E.-M., Gallouët, E., Baptist, F., Kuntz, M., & Cornic, G. (2005). Evidence for alternative electron sinks to photosynthetic carbon assimilation in the high mountain plant species *Ranunculus glacialis*. Plant, Cell & Environment, 28(9), 1123–1135. 10.1111/j.1365-3040.2005.01350.x

Takagi, D., Hashiguchi, M., Sejima, T., Makino, A., & Miyake, C. (2016). Photorespiration provides the chance of cyclic electron flow to operate for the redox-regulation of P700 in photosynthetic electron transport system of sunflower leaves. Photosynthesis Research, 129(3), 279–290. 10.1007/s11120-016-0267-5

Timm, S., Florian, A., Arrivault, S., Stitt, M., Fernie, A. R., & Bauwe, H. (2012). Glycine decarboxylase controls photosynthesis and plant growth. FEBS Letters, 586(20), 3692–3697. 10.1016/j.febslet.2012.08.027

Tiwari, R., Foyer, C. H., & Muscarella, R. (2026). Photorespiration as a thermal protection mechanism in trees: Measurements, assumptions, and persistent gaps. Global Change Biology Communications, 1(2), e70025. 10.1002/gcb4.70025

Tomeo, N. J., & Rosenthal, D. M. (2018). Photorespiration differs among *Arabidopsis thaliana* ecotypes and is correlated with photosynthesis. Journal of Experimental Botany, 69(21), 5191–5204. 10.1093/jxb/ery274

Trudeau, D. L., Edlich-Muth, C., Zarzycki, J., Scheffen, M., Goldsmith, M., Khersonsky, O., Avizemer, Z., Fleishman, S. J., Cotton, C. A. R., Erb, T. J., Tawfik, D. S., & Bar-Even, A. (2018). Design and in vitro realization of carbon-conserving photorespiration. Proceedings of the National Academy of Sciences of the United States of America, 115(49), E11455–E11464. 10.1073/pnas.1812605115

Ubierna, N., & Farquhar, G. D. (2014). Advances in measurements and models of photosynthetic carbon isotope discrimination in C3 plants: Photosynthetic carbon isotope discrimination in C3plants. Plant, Cell & Environment, 37(7), 1494–1498. 10.1111/pce.12346

Valentini, R., Epron, D., de Angelis, P., Matteucci, G., & Dreyer, E. (1995). *In situ* estimation of net CO_2_ assimilation, photosynthetic electron flow and photorespiration in Turkey oak (*Q. cerris* L.) leaves: diurnal cycles under different levels of water supply. Plant, Cell & Environment, 18(6), 631–640. 10.1111/j.1365-3040.1995.tb00564.x

van Bodegom, P. M., Douma, J. C., & Verheijen, L. M. (2014). A fully traits-based approach to modeling global vegetation distribution. Proceedings of the National Academy of Sciences of the United States of America, 111(38), 13733–13738. 10.1073/pnas.1304551110

Villar, R., Held, A. A., & Merino, J. (1995). Dark leaf respiration in light and darkness of an evergreen and a deciduous plant species. Plant Physiology, 107(2), 421–427. 10.1104/pp.107.2.421

Voss, I., Sunil, B., Scheibe, R., & Raghavendra, A. S. (2013). Emerging concept for the role of photorespiration as an important part of abiotic stress response. *Plant Biology (Stuttgart*, Germany), 15(4), 713–722. 10.1111/j.1438-8677.2012.00710.x

Wada, S., Miyake, C., Makino, A., & Suzuki, Y. (2020). Photorespiration coupled with CO_2_ assimilation protects photosystem I from photoinhibition under moderate poly(ethylene glycol)-induced osmotic stress in rice. Frontiers in Plant Science, 11, 1121. 10.3389/fpls.2020.01121

Walker, A. P., Johnson, A. L., Rogers, A., Anderson, J., Bridges, R. A., Fisher, R. A., Lu, D., Ricciuto, D. M., Serbin, S. P., & Ye, M. (2021). Multi-hypothesis comparison of Farquhar and Collatz photosynthesis models reveals the unexpected influence of empirical assumptions at leaf and global scales. Global Change Biology, 27(4), 804–822. 10.1111/gcb.15366

Walker, B. J., Orr, D. J., Carmo-Silva, E., Parry, M. A. J., Bernacchi, C. J., & Ort, D. R. (2017). Uncertainty in measurements of the photorespiratory CO_2_ compensation point and its impact on models of leaf photosynthesis. Photosynthesis Research, 132(3), 245–255. 10.1007/s11120-017-0369-8

Walker, B. J., VanLoocke, A., Bernacchi, C. J., & Ort, D. R. (2016). The Costs of Photorespiration to Food Production Now and in the Future. Annual Review of Plant Biology, 67(Volume 67, 2016), 107–129. 10.1146/annurev-arplant-043015-111709

Wingler, A., Lea, P. J., Quick, W. P., & Leegood, R. C. (2000). Photorespiration: metabolic pathways and their role in stress protection. *Philosophical Transactions of the Royal Society of London. Series B*, Biological Sciences, 355(1402), 1517–1529. 10.1098/rstb.2000.0712

Xu, Y., Fu, X., Sharkey, T. D., Shachar-Hill, Y., & Walker, A. B. J. (2021). The metabolic origins of non-photorespiratory CO_2_ release during photosynthesis: a metabolic flux analysis. Plant Physiology, 186(1), 297–314. 10.1093/plphys/kiab076

Ye, Z., Hu, W., Zhou, S., Robakowski, P., Kang, H., An, T., Wang, F., Xiao, Y., & Yang, X. (2025). Limitations of the Farquhar-von Caemmerer-berry model in estimating the maximum electron transport rate: Evidence from four C3 species. Biology, 14(6), 630. 10.3390/biology14060630

Ye, Z.-P., Liu, Y.-G., Kang, H.-J., Duan, H.-L., Chen, X.-M., & Zhou, S.-X. (2019). Comparing two measures of leaf photorespiration rate across a wide range of light intensities. Journal of Plant Physiology, 240(153002), 153002. 10.1016/j.jplph.2019.153002

Ye, Z.-P., Yang, X.-L., Ye, Z.-W.-Y., An, T., Duan, S.-H., Kang, H.-J., & Wang, F.-B. (2025). Evaluating photosynthetic models and their potency in assessing plant responses to changing oxygen concentrations: a comparative analysis of *A* _n_-*C* _a_ and *A* _n_-*C* _i_ curves in *Lolium perenne* and *Triticum aestivum*. Frontiers in Plant Science, 16, 1575217. 10.3389/fpls.2025.1575217

Yeo, M. E., Yeo, A. R., & Flowers, T. J. (1994). Photosynthesis and photorespiration in the genus *Oryza*. Journal of Experimental Botany, 45(5), 553–560. 10.1093/jxb/45.5.553

Yin, X., & Amthor, J. S. (2024). Estimating leaf day respiration from conventional gas exchange measurements. The New Phytologist, 241(1), 52–58. 10.1111/nph.19330

Yoshimura, Y., Kubota, F., & Hirao, K. (2001). Estimation of photorespiration rate by simultaneous measurements of CO_2_, gas exchange rate, and chlorophyll fluorescence quenching in the C_3_ plant Vigna radiata (L.) wilczek and the C_4_ plant *Amaranthus mongostanus* L. Photosynthetica, 39(3), 377–382. 10.1023/a:1015178209845

Zelitch, I., Schultes, N. P., Peterson, R. B., Brown, P., & Brutnell, T. P. (2009). High glycolate oxidase activity is required for survival of maize in normal air. Plant Physiology, 149(1), 195–204. 10.1104/pp.108.128439

Zhang, C., Zhan, D. X., Luo, H. H., Zhang, Y. L., & Zhang, W. F. (2016). Photorespiration and photoinhibition in the bracts of cotton under water stress. Photosynthetica, 54(1), 12–18. 10.1007/s11099-015-0139-9

Zhang, G., Zhang, L., & Wen, D. (2018). Photosynthesis of subtropical forest species from different successional status in relation to foliar nutrients and phosphorus fractions. Scientific Reports, 8(1), 10455. 10.1038/s41598-018-28800-4

Zhu, X.-G., Long, S. P., & Ort, D. R. (2010). Improving photosynthetic efficiency for greater yield. Annual Review of Plant Biology, 61(1), 235–261. 10.1146/annurev-arplant-042809-112206

Ziegler, C., Dusenge, M. E., Nyirambangutse, B., Zibera, E., Wallin, G., & Uddling, J. (2020). Contrasting dependencies of photosynthetic capacity on leaf nitrogen in early- and late-successional tropical Montane tree species. Frontiers in Plant Science, 11, 500479. 10.3389/fpls.2020.500479

