## Supplementary Section 3 for "Large differences in photorespiration and its temperature response among temperate trees"

### Supplementary Information

**Supplementary Table S1**: Data gathered from the extant literature on photorespiration rates estimated via some kind of gas exchange measure. Values plotted in Box 1.

| Study | Species | ϕ ± SE | $T_{leaf}$ (℃) |
|---|---|---|---|
| | $A_{net}$ based studies | | |
| Aliyev 2012 | Wheat (*Triticum aestivum* Azamatli-95) | 0.3 | 25 |
| Aliyev 2012 | Wheat (*Triticum aestivum* Giymatli-2/17) | 0.24 | 25 |
| Aliyev 2012 | Wheat (*Triticum aestivum* Gyrmyzy gul) | 0.3 | 25 |
| Bernacchi et al., 2001 | Tobacco (*Nicotiana tabacum*) | 0.271±0.015 | 25 |
| Cornic & Ghashghaie, 1991 | Common bean (*Phaseolus vulgaris*) | 0.44 | 25 |
| Keys et al., 1977 | Wheat (*Triticum aestivum*) | 0.114±0.024 | 20.5 |
| Oberhauer & Edwards 1993 | *Flaveria pringlei* | 0.924±0.097 | 25 |
| Sharkey 1985 | Common pea *(Pisum vulgaris)* | 0.352±0.04 | 25 |
| Takagi et al 2016 | Common sunflower (*Helianthus annuus*) | 0.998 | 25 |
| Tomeo & Rosenthal 2018 | *Arabidopsis thaliana* | 0.380±0.320 | 25 |
| Valentini et al., 1997 | Turkey oak, *Quercus cerris,* (Sept) | 0.622±0.047 | |
| Villalobos-Gonzalez et al 2022 | Grapevine, (*Vitis vinifera* chardonnay) | 0.828 | 28 |
| Villalobos-Gonzalez et al 2022 | Grapevine, (*Vitis vinifera cabernet sauvignon*) | 0.455 | 28 |
| Villalobos-Gonzalez et al 2022 | Grapevine, (*Vitis vinifera sauvignon blanc*) | 0.635 | 28 |
| Ye et al., 2019 | Wheat (*Triticum aestivum Z39-118*) | 0.398 | 30 |
| Ye et al., 2019 | Wheat (*Triticum aestivum Z39-118*) | 0.32 | 30 |
| Yeo et al 1994 | Rice (*Oryza sativa*) | 0.312 | 25 |
| Yeo et al 1994 | Rice (*Oryza australiensis*) | 0.372 | 25 |
| Yeo et al 1994 | Rice (*Oryza rufipogon*) | 0.263 | 25 |
| | ETR partitioning approach | | |
| Valentini et al., 1997 | Turkey oak (*Quercus cerris, sept*) | 0.827±0.079 | |
| Erel et al., 2015 | Olive (*Olea europaea*) | 0.30 | 25 |
| Hochberg et al 2013 | Grapevine, *Vitis vinifera* | 0.44±0.056 | 25 |
| Huang et al., 2015 | *Arabidopsis thaliana* | 0.43 | 25 |
| Zhang et al 2016 | Tree-liana (*Brideliya stipularis*) | 0.497 | 30 |
| Zhang et al 2016 | Tree-liana (*Strophioblachia fimbricalyx*) | 0.513 | 30 |

**Aliyev JA. 2012**. Photosynthesis, photorespiration and productivity of wheat and soybean genotypes. *Physiologia plantarum* **145**: 369–383.

**Bernacchi CJ, Singsaas EL, Pimentel C, Portis AR Jr, Long SP. 2001**. Improved temperature response functions for models of Rubisco-limited photosynthesis. *Plant, cell & environment* **24**: 253–259.

**Cornic G, Ghashghaie J. 1991**. Effect of temperature on net CO2 assimilation and photosystem II quantum yield of electron transfer of French bean (*Phaseolus vulgaris* L.) leaves during drought stress. *Planta* **185**: 255–260.

**Erel R, Yermiyahu U, Ben-Gal A, Dag A, Shapira O, Schwartz A. 2015**. Modification of non-stomatal limitation and photoprotection due to K and Na nutrition of olive trees. *Journal of plant physiology* **177**: 1–10.

**Hochberg U, Degu A, Fait A, Rachmilevitch S. 2013**. Near isohydric grapevine cultivar displays higher photosynthetic efficiency and photorespiration rates under drought stress as compared with near anisohydric grapevine cultivar. *Physiologia plantarum* **147**: 443–452.

**Huang W, Hu H, Zhang S-B. 2015**. Photorespiration plays an important role in the regulation of photosynthetic electron flow under fluctuating light in tobacco plants grown under full sunlight. *Frontiers in plant science* **6**.

**Keys AJ, Sampaio EVSB, Cornelius MJ, Bird IF. 1977**. Effect of temperature on photosynthesis and photorespiration of wheat leaves. *Journal of experimental botany* **28**: 525–533.

**Oberhuber W, Edwards G. 1993**. Temperature dependence of the linkage of quantum yield of photosystem II to CO2 fixation in C4 and C3 plants. *Plant physiology* **101**: 507–512.

**Sharkey TD. 1985**. O(2)-insensitive photosynthesis in c(3) plants : its occurrence and a possible explanation. Plant physiology **78**: 71–75.

**Takagi D, Hashiguchi M, Sejima T, Makino A, Miyake C. 2016**. Photorespiration provides the chance of cyclic electron flow to operate for the redox-regulation of P700 in photosynthetic electron transport system of sunflower leaves. *Photosynthesis research* **129**: 279–290.

**Tomeo NJ, Rosenthal DM. 2018**. Photorespiration differs among Arabidopsis thaliana ecotypes and is correlated with photosynthesis. *Journal of experimental botany* **69**: 5191–5204.

**Valentini R, Epron D, de Angelis P, Matteucci G, Dreyer E. 1995**. *In situ* estimation of net $CO^2$ assimilation, photosynthetic electron flow and photorespiration in Turkey oak (*Q. cerris* L.) leaves: diurnal cycles under different levels of water supply. *Plant, cell & environment* **18**: 631–640.

**Villalobos-González L, Alarcón N, Bastías R, Pérez C, Sanz R, Peña-Neira Á, Pastenes C. 2022**. Photoprotection is achieved by photorespiration and modification of the leaf incident light, and their extent is modulated by the stomatal sensitivity to water deficit in grapevines. *Plants* **11**: 1050.

**Ye Z-P, Liu Y-G, Kang H-J, Duan H-L, Chen X-M, Zhou S-X. 2019**. Comparing two measures of leaf photorespiration rate across a wide range of light intensities. *Journal of plant physiology* **240**: 153002.

**Yeo ME, Yeo AR, Flowers TJ. 1994**. Photosynthesis and photorespiration in the genus Oryza. *Journal of experimental botany* **45**: 553–560.

**Zhang SB, Zhang JL, Cao KF. 2016**. Differences in the photosynthetic efficiency and photorespiration of co-occurring Euphorbiaceae liana and tree in a Chinese savanna. *Photosynthetica* **54**: 438–445.

**Supplementary Table S2**: Mixed-effects model results for apparent photorespiration and photosynthesis parameters in response to species and leaf temperature

| Response variable | Fixed effects | (Random effects) | *F*-value | df1, df2 | *P*-value |
|---|---|---|---|---|---|
| $A_{net}$ | Species | (1 \| Replicates) | 18.19 | 6, 86 | **<0.001** |
| | $T_{leaf}$ | (1 \| Species) | 0.01 | 1, 115 | 0.92 |
| | Species × $T_{leaf}$ | (1 \| Replicates) | 1.38 | 6, 79 | 0.232 |
| $L_{app}$ | Species | (1 \| Replicates) | 5.73 | 6, 86 | **<0.001** |
| | $T_{leaf}$ | (1 \| Species) | 26.17 | 1, 115 | **<0.001** |
| | Species × $T_{leaf}$ | (1 \| Replicates) | 1.62 | 6, 79 | 0.15 |
| $\phi = L_{app} / A_{net}$ | Species | (1 \| Replicates) | 4.29 | 6, 86 | **<0.001** |
| | $T_{leaf}$ | (1 \| Species) | 17.63 | 1, 115 | **<0.001** |
| | Species × $T_{leaf}$ | (1 \| Replicates) | 0.90 | 6, 79 | 0.49 |

**Supplementary Table S3**: Temperature dependence of ETR-based photorespiration partitioning metrics.

| Metric | Definition | Slope (°C$^{-1}$) | Notes |
|---|---|---|---|
| $f_{Anet}$ | ETR to net assimilation / ambient ETR | − 0.0233 | Fraction of *ETR* to $A_{net}$ decreases with temperature; complementary to $f_{Loss}$ |
| $f_{Loss}$ | ETR to apparent loss / ambient ETR | + 0.0233 | Fraction of ETR to apparent loss increases with temperature |
| $R_p$, ETR | ETR-based apparent photorespiratory $CO_2$ loss | + 0.401 | µ mol e$^-$ m$^{-2}$ s$^{-1}$ per °C |
| $\phi$ *ETR* | $R_p$, ETR / $A_{net}$, ambient | + 0.0403 | Proportional apparent loss increases with temperature |

**Supplementary Table S4**: Photosynthesis ($A_{net}$) and photorespiration ($R_p$) rates in seven broadleaf tree species (Mean ± SE). numbers in parenthesis indicate the number of trees measured (biological replicates)

| | Photosynthesis rate ($A_{net}$) | | | Apparent loss of $CO_2$ assimilation rate ($L_{App}$) | | |
|---|---|---|---|---|---|---|
| | 25 °C | 30 °C | 35 °C | 25 °C | 30 °C | 35 °C |
| *Acer platanoides* (10) | 8.31 ± 1.22 | 9.39 ± 1.06 | 8.01 ± 1.36 | 1.62 ± 0.49 | 3.40 ± 0.54 | 4.13 ± 0.70 |
| *Betula pendula* (7) | 17.2 ± 1.30 | 14.3 ± 1.34 | 14.6 ± 1.19 | 2.78 ±1.58 | 6.97 ± 0.84 | 7.00 ± 0.58 |
| *Betula pubescens* (5) | 11.9 ± 1.59 | 10.2 ± 1.67 | 10.6 ± 1.89 | 4.43 ± 0.57 | 4.31 ± 0.77 | 4.13 ± 0.72 |
| *Corylus avellana* (7) | 8.25 ± 1.15 | 6.72 ± 0.71 | 5.82 ± 0.57 | 2.31 ± 0.60 | 3.51 ± 0.88 | 3.83 ± 0.48 |
| *Fagus sylvatica* (9) | 8.40 ± 0.98 | 12.5 ± 1.50 | 11.4 ± 1.07 | 2.25 ± 0.59 | 5.56 ± 1.10 | 7.86 ± 0.88 |
| *Scandosorbus intermedia* (8) | NA | 9.24 ± 0.73 | 10.0 ± 1.42 | NA | 6.33 ± 1.84 | 7.83 ± 1.74 |
| *Tilia cordata* (17) | 8.24 ± 0.40 | 10.0 ± 1.08 | 9.49 ± 1.11 | 2.90 ± 0.56 | 4.17 ± 0.56 | 4.65 ± 0.31 |

**Supplementary Table S5**: Apparent photorespiration to photosynthesis ratio ($\phi = L_{app} / A_{net}$) in seven broadleaf tree species (mean ± SE)

| | $\phi = L_{app} / A_{net}$ | | |
|---|---|---|---|
| | 25 °C | 30 °C | 35 °C |
| *Acer platanoides* | 0.22 ± 0.07 | 0.34 ± 0.06 | 0.52 ± 0.09 |
| *Betula pendula* | 0.18 ± 0.10 | 0.55 ± 0.12 | 0.48 ± 0.05 |
| *Betula pubescens* | 0.41 ± 0.02 | 0.42 ± 0.01 | 0.39 ± 0.03 |
| *Corylus avellana* | 0.34 ± 0.11 | 0.49± 0.08 | 0.67 ± 0.06 |
| *Fagus sylvatica* | 0.29 ± 0.11 | 0.43 ± 0.03 | 0.71 ± 0.06 |
| *Scandosorbus intermedia* | NA | 0.69 ± 0.22 | 0.94 ± 0.25 |
| *Tilia cordata* | 0.36 ± 0.08 | 0.43 ± 0.05 | 0.43 ± 0.04 |

**Supplementary Figure S1**: Histograms of daytime air temperature data (07:00 to 18:00, hourly intervals) for July, pooled from quality-checked SMHI station 97510 (Uppsala University; 59.8471°N, 17.6320°E, 23.5 m a.s.l.), covering 1985–2023.

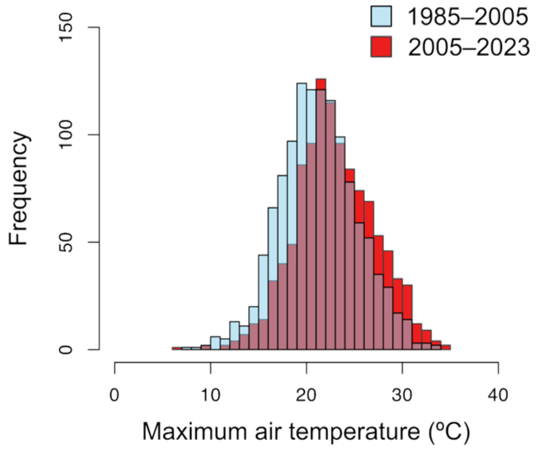


SMHI. (2024). Meteorological Observations from Station 97510 (Uppsala Aut) [Data set]. In *Swedish Meteorological and Hydrological Institute (SMHI).* https://www.smhi.se/data/meteorologi/ladda-ner-meteorologiska-observationer#param=airHumidity,stations=core,stationid=97510

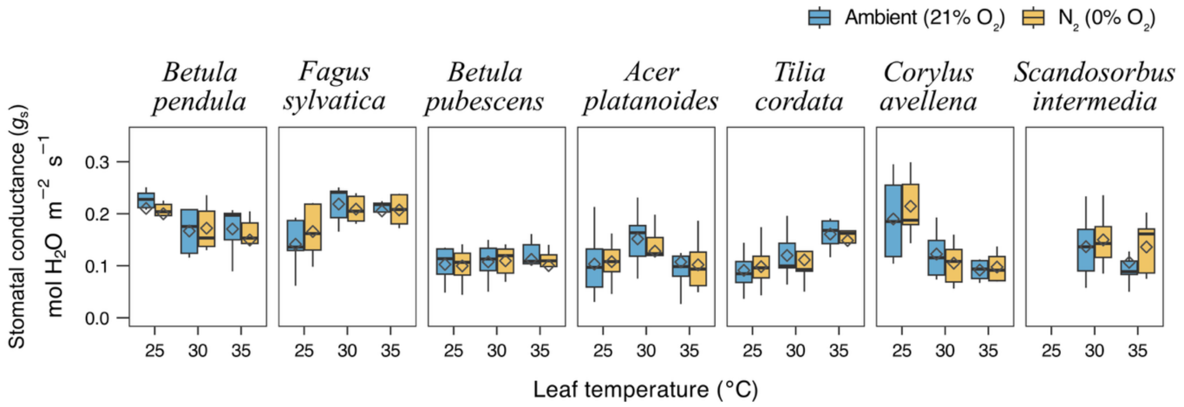


**Supplementary Figure S2**: Stomatal conductance ($g_s$) responses to ambient and $N_2$ conditions across species and temperatures. gs was measured for the same seven temperate broadleaf tree species at leaf temperatures of 25, 30, and 35 °C under ambient $O_2$ (21% $O_2$, blue) and $N_2$ (0% $O_2$, gold). Diamonds represent mean values. Across species and temperatures, gs differed only modestly between ambient and $N_2$, consistent with the absence of significant species or temperature effects in $\Delta\, g_s$ (all $P > 0.2$), suggesting that the $O_2$ shift primarily affected biochemical rather than stomatal processes.

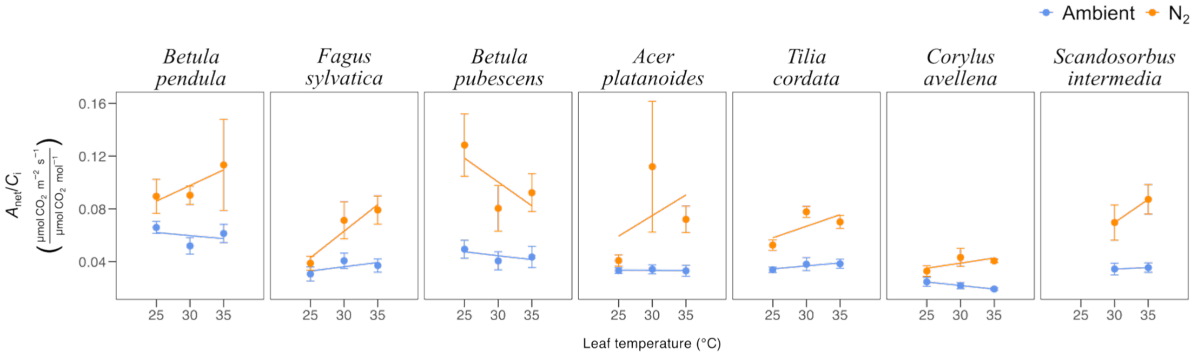


**Supplementary Figure S3**: Species-specific temperature responses of the ratio of $CO_2$ assimilation rate to intercellular $CO_2$ concentration ($A_{net}/C_i$) under ambient $O_2$ (21% $O_2$, blue) and $N_2$ (0% $O_2$, orange) at leaf temperatures of 25, 30, and 35 °C for seven temperate broadleaf tree species in Uppsala, Sweden; points show species means ± SE and lines represent linear fits across temperatures. Net photosynthesis per unit $C_i$ generally declined with temperature under ambient $O_2$, whereas suppression of photorespiration under $N_2$ tended to increase $A_{net}/C_i$ or maintain higher values at warm temperatures, indicating that photorespiration reduces the apparent efficiency of net $CO_2$ assimilation per unit internal $CO_2$ in these temperate tree leaves. In several species, the separation between ambient and $N_2$ $A_{net}/C_i$ was most pronounced at 35 °C, consistent with stronger photorespiratory limitation of net photosynthesis at high leaf temperatures.

**Supplementary Section N1: Temperature dependence of TPU-consistent $O_2$-shift responses**

Photorespiration is normally suppressed under a low-oxygen atmosphere because Rubisco's oxygenase activity requires $O_2$, and this suppression is expected to increase net $CO_2$ assimilation ($A_{net}$) by eliminating the competing oxygenation reaction. However, the resulting surge in carbon fixation can outpace the leaf's capacity to synthesise starch and export sucrose, a condition known as triose-phosphate-utilisation (TPU) limitation, in which inorganic phosphate becomes trapped in phosphorylated intermediates and the Calvin-Benson cycle is transiently constrained (Cornic & Louason, 1980; Sharkey & Vassey, 1989). Under TPU limitation, $A_{net}$ can therefore fail to increase, or even decline, when photorespiration is suppressed, despite the removal of the $O_2$-dependent $CO_2$ loss pathway.

Of 123 total ambient–$N_2$ measurement pairs, 24 (19.5%) showed a response pattern consistent with TPU limitation under the $N_2$ treatment ($L_{app} \leq 0$ or *ETR* loss $\leq 0$) and were excluded from ETR partitioning and related temperature-response analyses (see Methods). The incidence of TPU-consistent pairs declined sharply with increasing leaf temperature, from 34.8% at 25°C to 17.1% at 30°C and 5.6% at 35°C (binomial GLM: temperature effect $\chi^2_{(1)} = 14.64$, $P < 0.001$; species effect $\chi^2_{(6)} = 11.25$, $P = 0.081$), indicating that TPU-consistent suppression of net assimilation was most frequent at the cooler end of our measurement range and became progressively rare under heat-wave conditions. This pattern matches the established temperature sensitivity of TPU limitation, which is more likely to constrain photosynthesis at temperatures below a leaf's thermal optimum, when carbohydrate export and storage capacity lags behind Rubisco-driven carbon fixation (Stitt, 1986; Sage & Sharkey, 1987; Labate & Leegood, 1988; McClain & Sharkey, 2019). TPU-consistent pairs did not differ significantly from the remaining pairs in ambient net photosynthesis (Welch's $T_{(32.3)} = 1.27$, $P = 0.212$), ruling out low signal or measurement noise as the underlying cause and supporting a genuine physiological origin for this exclusion criterion.

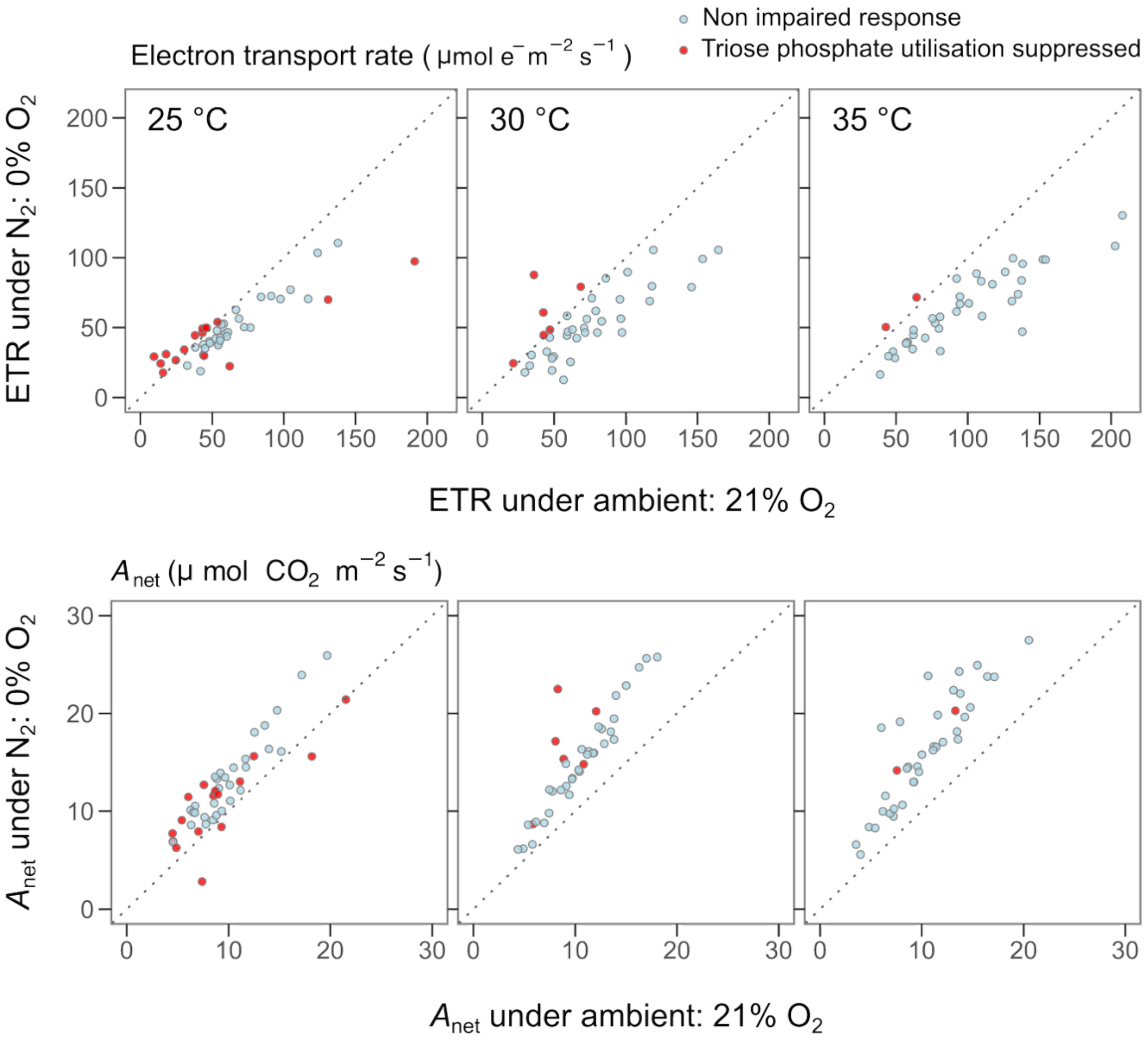

| $T_{\text{Leaf}}$ | Group | N | $\Delta A_{\text{net}}$ ($\mu$ mol $CO_2$ $m^{-2}$ $s^{-1}$) | $\Delta$ETR ($\mu$ mol $e^{-}$ $m^{-2}$ $s^{-1}$) |
|---|---|---|---|---|
| 25 °C | Unimpaired-response pairs | 30 | 3.1 ± 0.3 | –15.2 ± 1.7 |
| | TPU-suppressed pairs | 16 | 1.8 ± 0.7 | –8.4 ± 7.6 |
| 30 °C | Unimpaired-response pairs | 33 | 4.4 ± 0.3 | –24.7 ± 3.1 |
| | TPU-suppressed pairs | 6 | 7.5 ± 1.7 | 14.4 ± 7.9 |
| 35 °C | Unimpaired-response pairs | 36 | 5.8 ± 0.5 | –35.4 ± 3.6 |
| | TPU-suppressed pairs | 2 | 6.8 ± 0.2 | 7.4 ± 0.2 |


**Cornic G**, **Louason G**. **1980**. The effects of $O_2$ on net photosynthesis at low temperature (5°C). *Plant, Cell & Environment* **3**: 149–157.

**Labate CA**, **Leegood RC**. **1988**. Limitation of photosynthesis by changes in temperature : Factors affecting the response of carbon-dioxide assimilation to temperature in barley leaves: Factors affecting the response of carbon-dioxide assimilation to temperature in barley leaves. *Planta* **173**: 519–527.

**McClain AM**, **Sharkey TD**. **2019**. Triose phosphate utilization and beyond: from photosynthesis to end product synthesis. *Journal of Experimental Botany* **70**: 1755–1766.

**Sage RF**, **Sharkey TD**. **1987**. The effect of temperature on the occurrence of O(2) and CO(2) insensitive photosynthesis in field grown plants. *Plant Physiology* **84**: 658–664.

**Sharkey TD**, **Vassey TL**. **1989**. Low oxygen inhibition of photosynthesis is caused by inhibition of starch synthesis. *Plant Physiology* **90**: 385–387.

**Stitt M**. **1986**. Limitation of photosynthesis by carbon metabolism : I. evidence for excess electron transport capacity in leaves carrying out photosynthesis in saturating light and CO(2): I. Evidence for excess electron transport capacity in leaves carrying out photosynthesis in saturating light and CO2. *Plant Physiology* **81**: 1115–1122.
